# Broken force dispersal network in tip-links by the mutations induces hearing-loss

**DOI:** 10.1101/614610

**Authors:** Jagadish P. Hazra, Amin Sagar, Nisha Arora, Debadutta Deb, Simerpreet Kaur, Sabyasachi Rakshit

## Abstract

Tip-link as force-sensor in the hearing conveys the mechanical force originating from sound to ion-channels while maintaining the integrity of the entire sensory assembly in inner-ear. This delicate balance between structure and function of tip-links is regulated by Ca^2+^-ions present in endolymph. Mutations at the Ca^2+^-binding sites of tip-links often lead to congenital deafness, sometimes syndromic defects impairing vision along with hearing. Although such mutations are already identified, it is still not clear how the mutants alter the structure-function properties of the force-sensors associated with diseases. With an aim to decipher the differences in force-conveying properties of the force-sensors in molecular details, we identified the conformational variability of mutant and wild-type tip-links at the single-molecule level using FRET at the endolymphatic Ca^2+^ concentrations and subsequently measured the force-responsive behavior using single-molecule force spectroscopy with an AFM. AFM allowed us to mimic the high and wide range of force ramps (10^3^ - 10^6^ pN.s^−1^) as experienced in the inner ear. We performed *in silico* network analyses to learn that alterations in the conformations of the mutants interrupt the natural force-propagation paths through the sensors and make the mutant tip-links vulnerable to input forces from sound stimuli. We also demonstrated that a Ca^2+^ rich environment can restore the force-response of the mutant tip-links which may eventually facilitate the designing of better therapeutic strategies to the hearing loss.

**Significance Statement:** Force-sensors in inner ear are the key components in the hearing. Mutations in force-sensors often lead to congenital hearing loss. Loss of hearing has become a threat to humanity, with over 5% of world population suffering from deafness and 40% of which is congenital, primarily due to mutations in the sensory machinery in inner-ear. A better understanding of the molecular mechanism of the underlined hearing loss due to mutations is, therefore, necessary for better therapeutics to deaf. Here with a zoomed region of the force-sensors, we pointed out the differences in the force-propagation properties of the mutant and wild-type force-sensors. Our observation on restoring of functions of mutants in Ca^2+^-rich buffer indicates methods of developing low-cost therapeutic strategies against deafness.

## Introduction

Mechanical force is transduced into an electrical signal in hearing (1). Tip-links, a proteinaceous bridge between adjacent stereocilia of hair-cells (2–4), receives the input-force and as a gating-spring in the mechanotransduction in the hearing conveys the force to mechanosensory ion-channels (5). Molecular components that orchestrate tip-links are two non-classical cadherins, cadherin-23(Cdh23) and protocadherin-15(Pcdh15) (6–10). These proteins from the same stereocilium are engaged in parallel to form cis-homodimers through their extracellular (EC) regions (6, 11). The EC regions of the cadherins are made up of multiple domains arranged in tandem with inter-domain linkers. The two terminal domains (EC1-2) of the cis-homodimers from the opposing stereocilia interact in a “handshake” manner to form the trans-heteromeric tip-link complex.

The inter-domain linker regions of the cadherins are enriched with charged residues, including aspartate, asparagine that is conserved with Ca^2+^ binding motifs, such as DRX, DXNDN (12–16). Ca^2+^ ions in the environment bind to the motifs at the linkers and tune the elasticity of cadherins (17–20), thus the entropic conformations of the proteins in tip-links (21). The flexibility of the tip-links is also tuned by the molecular motors at low-Ca^2+^ (22). Reportedly 80% of mutations at the inter-domain Ca^2+^-binding linker regions are linked with deafness in humans and mice. These mutations alter the binding-affinity of the proteins to Ca^2+^-ions and directly or indirectly alter the structure-function equilibrium of the tip-links. Mutations at the linkers between domains (e.g., EC2-EC3 in Pcdh15) that are not engaged in trans-interactions are implicated to impaired cis-dimerization (11). Our interests lie on the mutations at the linkers of EC1-2 that reside at the binding interface but do not contribute in the trans-binding or cis-binding (11, 23–28). Two of such mutants are: Aspartate101 to Glycine in Cdh23 EC1-2(D101G) that is associated with non-syndromic deafness DFNB12 (26) and Aspartate157 to Glycine in Pcdh15 EC1-2(D157G) (Figure 1a) directly associated with syndromic hearing-loss diseases (USH1F) (29). The individual domains of Cdh23 EC1-2 wild-type (WT) (PDB ID: 2WCP) and mutant Cdh23 EC1-2(D101G) (PDB ID: 2WD0) are architecturally very similar too (Figure 1b) except with a subtle difference in the inter-domain arrangements (30). However, both the mutants have significantly lower Ca^2+^ binding affinity than the WT. The dissociation constants (*K_D_*) of four Ca^2+^ binding sites of Cdh23 EC1-2(WT) are 1.9, 71.4, 5.0, and 44.3 μM for sites 0 to 3 assigned sequentially from N-terminal (30). The *K_D_*s for site 1 and 3 might be interchanged as the assignment was merely a prediction based on the sequence similarities. Mutation at D101 which directly forms a complex with a Ca^2+^ ion at site 2, reduces the affinities cooperatively for Cdh23 EC1-2(D101G) with *K_D_* values measured as 3.9, 40.6, and >100μM (Table S1, Figure S1a) (30). For Pcdh15 EC1-2(WT), the *K_D_* values for three binding sites are 11.2, 34.6, and 42.5 μM in the ascending order whereas the same for Pcdh15 EC1-2(D157G) is 16.8, 44.3, and 152.7μM (Table S2, Figure S1b).

**Figure 1:**
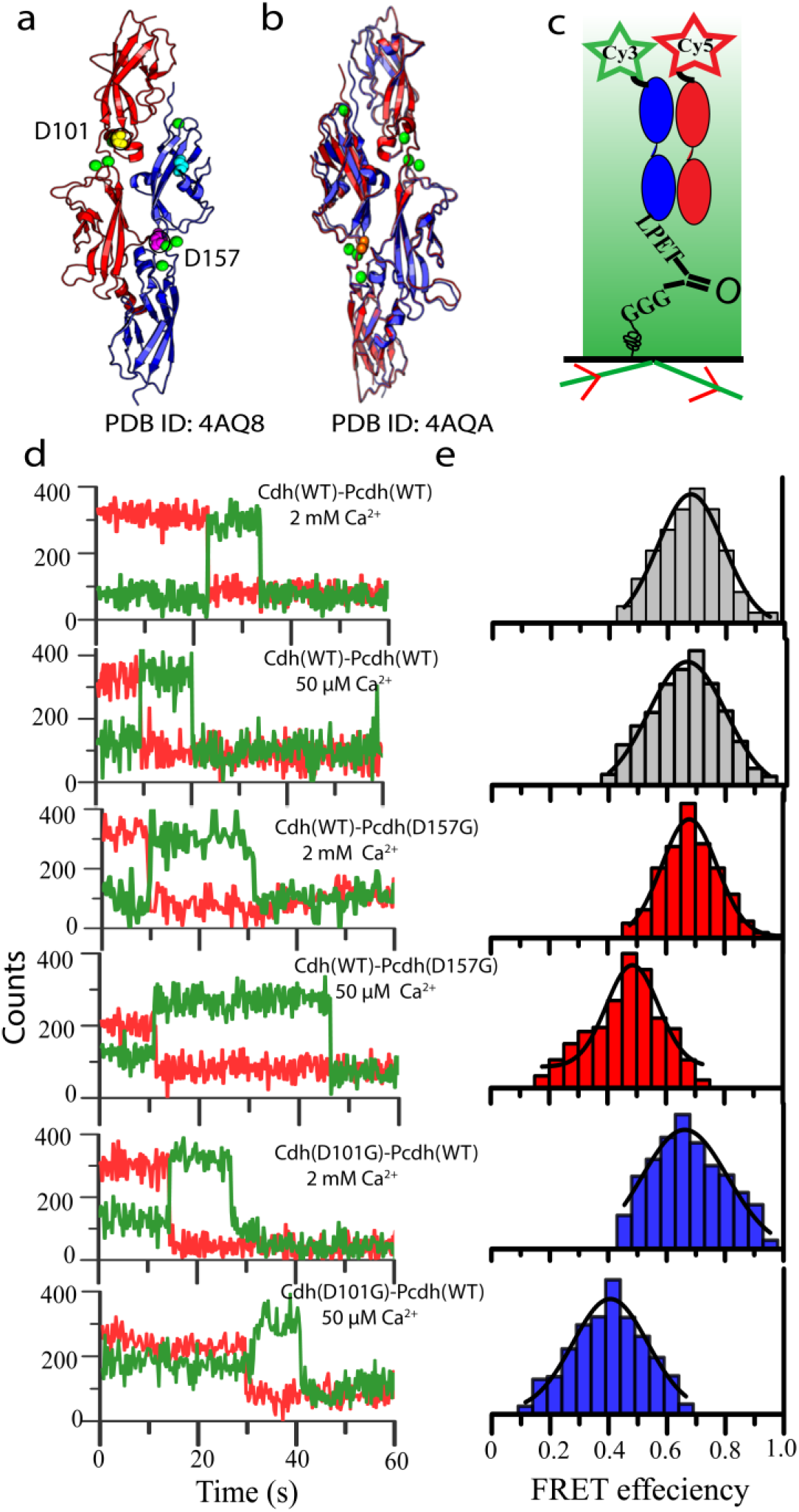
smFRET identifies different conformations for mutants in tip-links at low Ca^2+^: (a) Ribbon diagram of the wild-type tip-link highlighting the spatial locations of mutant, D101 for Cdh23 (Blue) and D157 for Pcdh15 (Red), and the calcium ions (orange) in the two outermost domains of the complex. (b) A representative structure of the mutant tip-link complex (Cdh23 (D101G) (blue) – Pcdh15 (WT)(red)) superimposed with the wild-type complex (Cyan) to highlight the structural differences as observed in Crystals. (c) Cartoon representation of the smFRET using a TIRF microscope. (d) Representative time traces of the fluorescence intensities from the donor (green) and acceptor (red) dyes for WT and mutant complexes at varying calcium concentrations. (e) The corresponding distributions of the FRET-efficiency are plotted for WT and mutant complexes at varying calcium concentrations.

The Ca^2+^-ion concentration in scala media of endolymph where tip-links are bathed varies from 20-60 μM along the tonotopic axis from the base to apex with a local flux of ∼50 μM around hair-bundles (31, 32). It is apparent, therefore, to speculate that with reduced Ca^2+^- affinity, both the mutants, Cdh23 EC1-2(D101G) and Pcdh15 EC1-2(D157G), will have lower Ca^2+^-occupancy compared to the WT in endolymph. Here, using single-molecule Forster Resonance Energy Transfer (smFRET) at the physiologically relevant Ca^2+^ (50 μM), we first verified whether mutants are able to form the tip-links and if so, what are the conformational differences between WT and mutant complexes. Tip-links, as force-conveyor, are exposed to mechanical force of various intensities periodically from the sound-stimuli. Using single-molecule force spectroscopy (SMFS), we exerted force ramps to the single tip-link complexes according to the tonotopic-map, measured the dissociation kinetics of the complexes, and deciphered how the parameters are altered in mutants. Finally, we proposed a general molecular mechanism for the differences in the force-resilience properties between all linker-specific mutants and WT, using the dynamic network analysis of force propagation *in silico*. Since only EC1-2 domains were used for all experiments, we avoid writing domain numbers with the protein variants.

## Results and Discussion

### Mutants in tip-links attain different conformations than WT at low Ca^2+^

We performed smFRET on a glass-coverslip using a Total Internal Reflection Fluorescence Microscopy (TIRFM) between Cdh23(WT) - Pcdh15(WT) (WT complex), Cdh23(WT) - Pcdh15(D157G) (mutant-1 complex), and Cdh23(D101G) - Pcdh15(WT) (mutant-2 complex) at 50 μM (low) of Ca^2+^ to mimic the physiological endolymph environment, and at 2 mM (high) of Ca^2+^ to saturate all the binding sites and replicate the conformations in the crystal structure. To maintain the tip-link orientation in our experiments, we always anchored the C-termini of the proteins specifically and covalently to surfaces (Figure 1a-c). For smFRET, we attached the C-termini of Cdh23 variants to coverslips using well-established sortagging chemistry (33). The N-termini of Cdh23 variants were recombinantly modified with cysteine at Valine (V2C) as Cdh23 did not contain any intrinsic cysteine and labeled the V2C site with Cy3 (donor, λ_ex_ = 545 nm; λ_em_= 563 nm) using thiol-maleimide Michael addition. The C-termini of Pcdh15 variants were labeled with Cy5 (acceptor, λ_ex_ = 645 nm; λ_em_= 670 nm) again using sortagging chemistry (Figure 1c and d, S4). We used iSMS (the Birkedal lab, Aarhus Universitat) for smFRET analysis (Materials and Methods, Figure 1d and S4) (34) and estimated the FRET-efficiency (*E_FRET_*) from the intensity ratios between acceptor and donor from the colocalized spots (Figure 1d). We used the peak-maxima of the distributions as most-probable efficiency 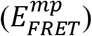.

A little difference in the distributions of *E_FRET_* for WT complex were recorded at high (2mM) and low (50 μM) Ca^2+^, and for both conditions the estimated 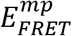 are 0.68±0.05 and 0.67±0.04 respectively (Figure 1d and e). The value of 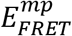 is relatively high and indicates the proximity of the dyes in the complex with a donor-acceptor distance (*d_D–A_*) *of* 4.62 nm ± 0.02 *nm*. No alteration in the distances possibly suggests a negligible change in the conformation of the complexes at two extreme Ca^2+^- concentrations. The distributions of *E_FRET_* for the mutant-complexes overlapped with the WT too, when recorded at high Ca^2+^ (2mM) (Figure 1d and e). This was expected from the crystal structure as well. However, at low Ca^2+^-concentration, we observed distinct efficiency-distributions for mutant-complexes with 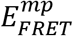 of 0.48±0.02 (*d_D–A_* = 5.26 ± 0.06 *nm*) and 0.40 ± 0.03 (*d_D–A_* = 5.56 ± 0.08 *nm*) for mutant-1 and mutant-2, respectively. FRET efficiency and distances in various complexes have been summarized in Table S3. For both the mutant-complexes, the distributions are shifted towards lower FRET efficiency than the WT complex, suggesting of different conformations for mutants at low Ca^2+^ with wider separations between donor-acceptor than WT. A more bend conformation for Cdh23 (D101G) than Cdh23(WT) at low-Ca^2+^ was already predicted from MD simulations alongwith wider distributions of the mutants in the conformational space (30). The MD simulations were done individually with the mutant proteins at unsaturated Ca^2+^-environment. However, we observed single conformations for both the mutants and WT tip-links within our bin-time of measurements. Similarities in the FWHM (Full Width at Half-Maxima) of the *E_FRET_* distributions for all constructs at varying Ca^2+^ also refer the same conformational variability in WT and mutant complexes. This anomaly between experiment and MD might be due to the differences in time-scales of measurements, as experimental output are time-averaged in milli-seconds whereas MD results are in pico-seconds.

Interestingly, both mutant complexes registered indistinguishable *d_D–A_*, irrespective of their mutation sites in different proteins in the complexes. For both the mutant-complexes, the *d_D–A_* measures the distance between Cdh23 EC1-2 N-terminal to Pcdh15 EC1-2 C-terminal (*d_NC_*), and this separation is estimated to be nearly equal irrespective of the spatial locations of the mutations. The indistinguishable *d_D–A_* for both the mutant-complexes suggests an equal extent of bent of the mutant proteins at 50μM Ca^2+^, provided the binding interface for both the mutant-complexes are identical. An increase in *d_D–A_* from the WT complex indicates the bent for the mutant protein is away from the WT counterpart in both the mutant complexes.

### Conformational differences in mutants alter the force-induced dissociation pathways

To decipher how the divergence in conformations alter the force-responsive behavior of the mutant complexes than WT, we performed single-molecule force-ramp spectroscopy (SMFS) using Atomic Force Microscope (AFM) at high (2mM) and low (50 μM) Ca^2+^ (Figure 2a). Use of AFM allows us to apply force ramps varying from 10^3^ pN.s^−1^ to 10^6^ pN.s^−1^, as in the tonotopic axis of the cochlea (35). Soft cantilevers with the stiffness of ∼20pN.nm^−1^, comparable to the stiffness of the hair-bundles, were used for the experiments (35). We performed SMFS for three different protein-pairs: Cdh23(WT) - Pcdh15(WT), Cdh23(D101G) - Pcdh15(WT), and Cdh23(WT) - Pcdh15(D157G). To maintain the native pulling geometry of tip-links, we covalently attached the C-termini of the interacting proteins to the cantilevers and glass coverslips (e.g., Cdh23 on cantilever against Pcdh15 on glass-coverslip or vice-versa) with Polyethylene glycol (PEG, molecular weight 5kDa) as the spacer. The covalent attachment was achieved using sortagging as described before (Figure 2a) (33). As a signature for specific protein-protein interactions, the force-stretching of PEG was monitored and fitted to Freely-Joint Chain model (FJC) (Equation 2 in Materials and Methods, Figure S5). On average, 5% of the total collected data showed such single-molecule force-stretching events (Figures S6, S7 and Table S4), which we fit with the FJC model. We estimated the contour-length (*lc*) of the PEG from the fit as 55 ±6 nm (Figure S7b). The *l_c_*-distribution corroborates with the theoretical estimation of *lc* obtained from the extension of two 5kDa PEG molecules from both the opposing surfaces aligned in series (36). Further, to confirm whether the single-molecule stretching events were resulting from specific protein-protein interactions, we repeated the SMFS in the absence of Ca^2+^ and expectedly, observed a significant drop (5% to 0.6%) in the frequency of the single-molecule stretching events. For each protein-pair, we performed SMFS at six different pulling velocities and calculated loading-rates (LR) for every force-extension curve using equation 3 (37). The unbinding forces (*F_ub_*) were measured from the peak of the force-stretching curves (Figure S6) and plotted as *F_ub_*-distribution for each velocity (Figure 2b and 2c). We fit each *F_ub_*-distribution to single Gaussian distribution and obtained the most probable unbinding forces (*F_mp_*) from the peak-maxima. Similarly, we also obtained the most-probable LR (*LR_mp_*) from distributions (Table S6). For the comparison of the unbinding forces, we overlaid the *F_ub_*-distributions of WT-pair with mutant-complexes (mutant-1 and mutant-2) for corresponding velocities. Experiments at high-Ca^2+^, we observed no notable differences in the force-distributions (Figure S8). The similarities in the force-distributions are corroborating with the smFRET results. However, the force-distributions were significantly different in low-Ca^2+^. We observed lower *F_ub_* for mutants than WT (Figure 2b). For all cases, *F_mp_* increased linearly with *LR_mp_* (Figure 2c). However, the extent of increment with increasing LR was greater for WT-pair than mutants. Mutants featured nearly identical *F_mp_* vs *LR_mp_* behavior with a distinctly different intercept than the WT. The monotonous increase in *F_mp_* with *LR_mp_* replicates the typical force-induced rupture behavior described in Bell-Evan’s model. This model assumes that the irreversible rupture of bonds induced by time-dependent external forces follow a free energy barrier-crossing quasi-adiabatically, where the distance between bound state to the transition state (*x_β_*) in the escape-potentials remains untampered to a small force. We fit the curves individually to Bell-Evan’s model (38) (Equation 4) and measured the intrinsic off-rate (*k^0^off*) and *x_β_* (Table 1) for all three pairs. Mutant complexes showed relatively faster dissociation than WT-complex (Figure 2c), however, the difference was not remarkable. We observed a similar trend in the intrinsic off-rates when measured in ensemble experiments using Bio-layer interferometry (BLI) (Figure S9 and Table S5). In fact, a significant difference in off-rate was not expected as the mutants influence the natural binding interface marginally. However, the magnitude of *x_β_* was more than double for both the mutant complexes (0.41±0.04 for mutant-1 and 0.44±0.04 for mutant-2) than WT(0.18 ± 0.03) (Figure 2d and Table 1). Interestingly higher *x_β_* employs higher external energy (*E_ext_* = *Fx_β_* cos *θ*) to tilt the escape-barrier towards dissociation, at fixed mechanical-tension. In other words, the extent of tilting of the escape-energy barrier towards dissociation is more for mutants than WT at same external forces. Higher tilting facilitates the dissociation of mutants manifolds faster than WT in presence of an external force 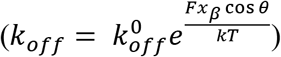. Mutants thus become more susceptible to detach in external mechanical-stimuli than WT complex.

**Figure 2:**
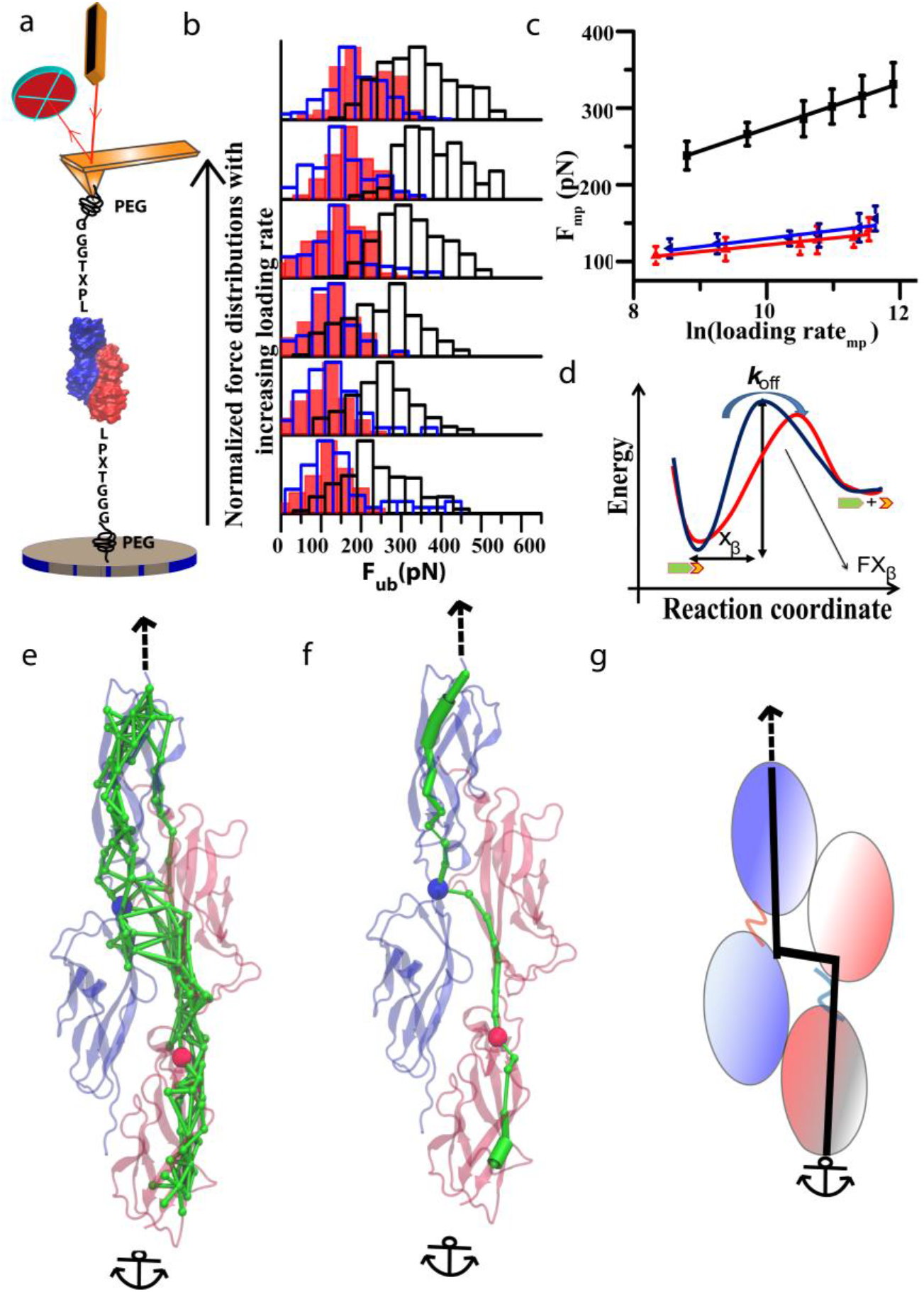
Mutants at low-Ca^2+^ follow different force-induced dissociation pathways than WT tip-links. (a) Cartoon representation of the single-molecule force spectroscopy setup using an AFM. (b) Distributions of unbinding forces obtained at different loading-rates are plotted for Cdh23(WT)–Pcdh15(WT) complex (black boundary, no shading), Cdh23(D101G)–Pcdh15(WT) complex (blue boundary and blue strip), Cdh23(WT)– Pcdh15(D157G) complex (red shaded). (c) The most probable unbinding forces (*F_mp_*) obtained at different loading rates (R) are plotted for Cdh23(WT)–Pcdh15(WT) complex (black), Cdh23(D101G)–Pcdh15(WT) complex (blue), Cdh23(WT)–Pcdh15(D157G) complex (red) with the natural log of loading rates. The corresponding fits to the Bell-Evan’s model are shown in solid lines with matching colors. (d) The force-induced dissociations of the complexes, the wild-type complex (blue) and mutant complex (red), as obtained from the data-fitting to Bell-Evan’s model in (c) are represented in a potential-energy diagram. (e) The sub-optimal force-propagation paths (green ribbons) obtained from the SMD of the WT-complex are overlaid on the ribbon-structure of the WT complex. D101 of Cdh23 and D157 of Pcdh15 are present on the suboptimal paths and shown as blue and red sphere respectively. The corresponding pulling directions are shown in arrow and anchor. (f) The most probable force-propagation, known as the optimal path of force-propagation, is shown separately on the complex in green ribbon. The presence of D101 of Cdh23 and D157 of Pcdh15 on the optimal path has been shown as blue and red spheres respectively. (g) The same optimal force-propagation is shown again in the cartoon model to highlight the distribution in force in the orthogonal direction to the force-pulling at the interface of the complex.

### The physiological significance of x_β_

*x_β_* has been extensively correlated with a wide variety of factors in literature to define the underlying escape-energy landscapes, frequently for the force-induced rupture events (39). It has been used to understand the conformational properties like ligand-induced changes in conformational variability in LacY (40), stiffness of polySUMO protein (41), as well as physical distances, for example, the stretching distance required to unfold Per-ARNT-Sim domain (42), depth of glucose in the translocation pathway of Sodium Glucose Co-transporter SGLT (43), and size of single unfolding unit in spectrin (44). *x_β_* in the current study measures the force-resistance or force-dissemination through the complexes, with higher the value of *x_β_* lesser is the resistance to dissociation under force. With an attempt to describe the force-dissemination pathway in the complexes with molecular details, we performed constant-velocity SMD at various pulling rates using QwikMD (45), a plugin of VMD (46) and run using NAMD (47, 48) (version 2.12). To mimic the geometry of SMFS experiments, we anchored the C-termini of Pcdh15 EC1-2(WT) and pulled from the C-termini of Cdh23 EC1-2(WT) or vice-versa, at four different pulling velocities (1 Å/ns, 2.5 Å/ns, 5 Å/ns and 7.5 Å/ns). Next, we performed the dynamic network analysis using VMD to identify the residues that move in concert with each other and determine the optimal path in the complex through which the external force propagates along the complex most frequently using Networkview plugin of VMD (49, 50)(Methods). Figure 2e depicts the suboptimal paths whereas, Figure 2f features only the optimal path. From the suboptimal paths, we observed that the force is transmitted from one protomer to the interacting partner at multiple points throughout the interface (Figure 2e). Multiple such intermolecular paths imply that the applied load is dispersed well at the binding-interface, thus reducing the probability of force-induced dissociation of the complex. Detailed analysis of the optimal paths revealed two further exciting points. Firstly, D101 of Cdh23 and D157 of Pcdh15 are on the optimal force-bearing path (Figure 2f). Mutations on the optimal path, if associated with structural-alterations, have been shown to steer the force-propagation differently than WT in the Dockerin-Cohesin complexes (51, 52). Accordingly, mutations in D101 and D157 are also expected to divert the force-propagation in the complexes as mutations in these residues attained bent-conformations at low-Ca^2+^ than WT. Secondly, we observed an orthogonal component in the optimal path aligning perpendicularly to the direction of pulling, and the position of this component is nearly at the center of the binding interface (Figure 2f, g). An orthogonal component effectively reduces the magnitude of the force exerting on the complex for dissociation, thus making it more resilient to force. Similar force-dissipation components orthogonal to the pulling direction were previously illustrated for silk crystalline units (53) and the ultrastable protein complex of X-module of Dockerin with Cohesin (51). In both cases, the orthogonal component was spatially located at the center of the interface of the domains or proteins as seen here (Figure 2f, g). It was also demonstrated that mutations on the optimal force-propagation path near the interface, not only alter the force-propagation but also diminish the orthogonal-component, thus leaving the complex more susceptible to force.

To elucidate the force-propagations in mutant complexes, we extended our in-silico measurements with mutants. However, the critical point to remember here is that we do not have definite knowledge on the occupancy of Ca^2+^ ions in mutant complexes at low-Ca^2+^ endolymph, and neither we have the corresponding crystal structures. Therefore, to mimic the structure of Cdh23 (WT) - PCdh15 (D157G) complex (mutant-1) at low calcium, we started with the crystal structure of the WT complex (PDB ID: 4AQ8), mutated Aspartate(D) at 157^th^ position of Pcdh15 to Glycine(G) using QwikMD (a tool in VMD), removed the calcium (at position 3, Figure S1b) interacting to D157, and then performed Gaussian Accelerated MD (GAMD)for 200 ns in triplicate(Methods). For mutant-2 complex, we began with the crystal structure of Cdh23 (D101G) – Pcdh15 (WT) complex (PDB ID: 4AQA) solved at high-Ca^2+^, removed the 2^nd^ calcium (interacting to D101 in the WT version, Figure S1a), and then performed GAMD as before. Both the complexes showed higher flexibility with higher fluctuations (root-mean-square fluctuations) at the linkers (Figure 3a-e). Detailed studies from the Cartesian Principal Component Analysis (cPCA) based cluster-analysis (using grcarma) (54) of the trajectories featured an emerging conformation for both the complexes with time (Figure S10b for D157G complex and Figure S10c for D101G complex). We, therefore, measured the interdomain angles of the mutant proteins throughout the simulation-time at an interval of 20 ps and plotted as distributions. To emphasize the change in interdomain angles, we plotted the angle-distribution for Pcdh15 (D157G) along with Pcdh15 (WT) for mutant-1 (Figure 3f) and Cdh23 (D101G) along with Cdh23 (WT) for mutant-2 complexes (Figure 3g). The bending in conformations of the mutant proteins in the complexes was also expected from the smFRET studies at low-Ca^2+^. Interestingly, *H-bond* analysis showed no alterations in the interacting residues for both the mutant complexes within the simulation time (Figure S11).

**Figure 3:**
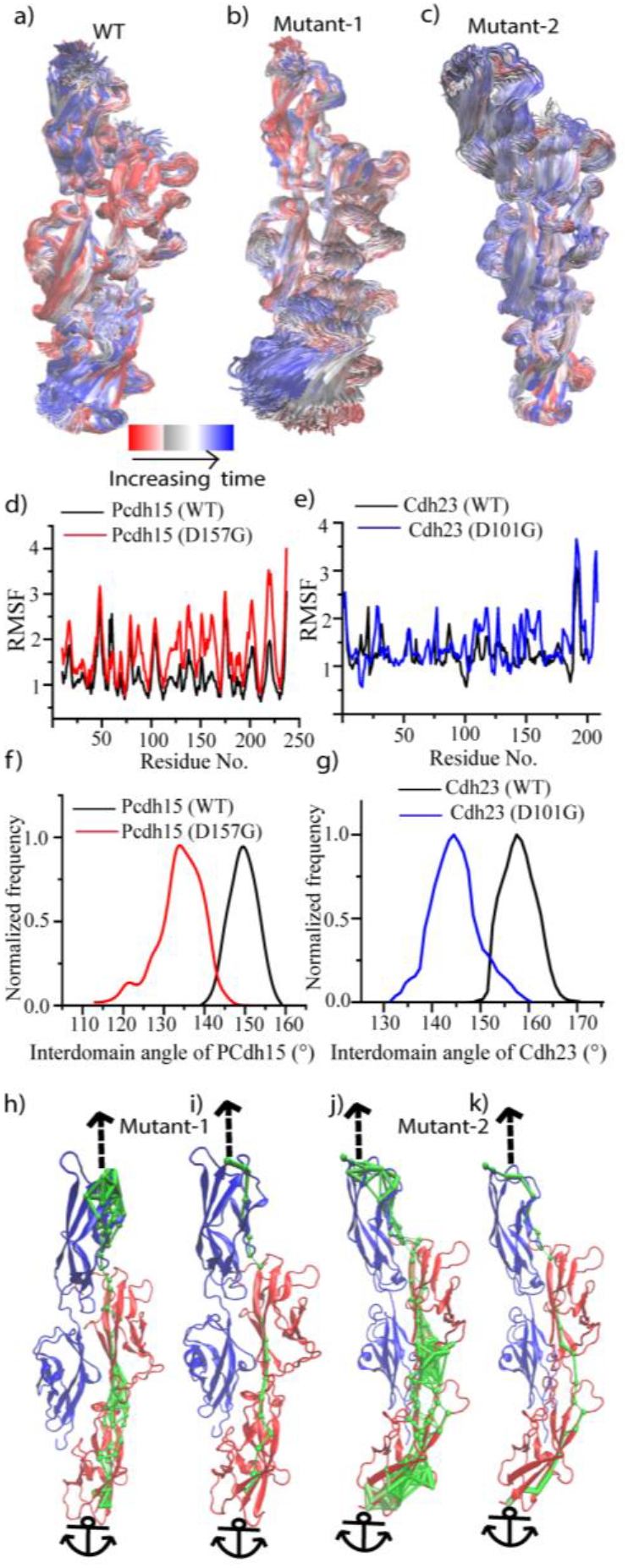
Mutant-complexes follow different and narrow force-dispersal paths with the broken orthogonal component as seen in the WT complex. Frames obtained after every 0.5 ns during 200 ns GAMD simulations for (a) Cdh23 (WT)–Pcdh15 (WT) complex with all-Ca^2+^, (b) Cdh23 (WT) – Pcdh15 (D157G) complex (mutant-1) after removing 3^rd^ Ca^2+^ from Pcdh15 (D157G), and (c) Cdh23 (D101G) – Pcdh15 (WT) complex (mutant-2) after removing 2^nd^ Ca^2+^ from Cdh23 (D101G). (d) Root-mean-square-fluctuations (RMSF) of the residues of Pcdh15 (WT) and Pcdh15 (D157) are compared for the simulations done in complex with Cdh23 (WT) as mentioned in (b). (e) RMSF of all the residues of Cdh23(WT) and Cdh23 (D101G) are compared for simulations mentioned in (c), performed in complex with Pcdh15 (WT). Both cases, mutants exhibit more fluctuation, notably at the linker regions, than WT entailing higher flexibility of the mutants at low Ca^2+^-environment. (f) Distribution of the angle between EC1 and EC2 domains of Pcdh15 (WT) and Pcdh15 (D157G) for the simulations described in (a), with Cdh23 (WT) with all-Ca^2+^ (black), and (b) with Cdh23 (WT) but 3^rd^-Ca^2+^ removed (red) respectively. The graph features a wider angle-distribution for the mutant than WT, with a lowering in peak-maxima to 134° for the mutant from 152° of the WT. (g) Distribution of the angle between EC1 and EC2 domains of Cdh23 (WT) and Cdh23 (D157G) for the simulations described in (a), with Pcdh15 (WT) with all-Ca^2+^ (black), and (c) with Pcdh15 (WT) but 2^nd^-Ca^2+^ removed from Cdh23 (red), respectively. The graph features a wider angle-distribution for the mutant than WT, with a lowering in peak-maxima to 142° for the mutant from 161° of the WT. (h) All the suboptimal force-propagation paths for Cdh23 (WT) – Pcdh15 (D157G) are overlaid on the ribbon-structure of the complex. (i) The optimal force-propagation path is shown for mutant-1. the (j) All the suboptimal force-propagation paths are shown in green for Cdh23 (D101G) – Pcdh15 (WT) complex. (k) The optimal force-propagation path is shown for mutant-2. Optimal paths for both the mutant complexes exhibit vanishing of perpendicular force dispersal path which was prominent in the WT complex.

We used the most populated bent conformations of the mutant complexes for SMD at constant velocities as discussed previously for the WT complex (Figure S12a and S12b) and performed dynamic network analysis to decipher the most-traveled force-propagation paths. To our expectations, the paths in both the mutant complexes are not only different than the WT complex, the orthogonal component that served as additional force-dissipation component for the WT complex also disappeared (Figure 3h, i for mutant-1 complex and figure 3j, k for mutant-2, and Figure S13a-c). The critical nodes and edges that help in force-dissipation in the WT-complex became fewer and narrower in both the mutant complexes (Figure S14a-c). Furthermore, all the suboptimal paths in the mutant complexes passed through one end of the binding-interface with an optimal path transversing from one protein to the partner at the initiation-site of the interface, not at the center of the interface as WT. This anisotropy in force-dispersion may impart the load on one bond at a time, similar to unzipping which registers low resistance to force.

## Conclusion

Congenital hearing loss is one of the predominant chronic symptoms in newborns from all over the world. Impaired hearing in babies is also associated with the delay in speech, language, emotional, and social developments. Mutations in the force-sensing assembly are considered as one of the major causes of congenital deafness in developed countries. Unfortunately, not much therapeutic strategies are known congenital deafness. Use of personalized hearing aids suffices the purpose marginally. A better understanding of the molecular mechanism of the underlined hearing loss due to mutations is, therefore, necessary to cater a better therapeutic solution to deaf newborns.

Ca^2+^ ions play a crucial role in maintaining the stiffness of the tip-links at low-Ca^2+^. Extensive studies have been done on the quantitative estimation of Ca^2+^ ions in endolymph and identifying the key molecular players that regulate the bulk, as well as the local endolymphatic Ca^2+^ concentration. However, the molecular mechanism of the altered force-conveying by mutant tip-links associated with severe deafness was not understood. Monitoring the conformational variations using smFRET at the endolymphatic Ca^2+^, we identified bent-conformations for mutants with a wider separation between two opposite termini (C- and N-) of the heterodimer. The wider separation is reflected as higher *x_β_* from our SMFS, measured at force ramps as experienced by tip-links in tonotopic axis at low-Ca^2+^. Accordingly, our *in-silico* network analysis of force-propagation indicated a difference in force-steering for mutants than WT and measured shorter force-dissipation paths for mutants. Shorter force-dissipation paths facilitate a significantly faster dissociation for the mutants than WT under force from sound-stimuli. Surprisingly, mutants featured identical force-conveying to WT at high-Ca^2+^. Such distinct differences in the force-dissemination by the WT and mutant tip-links, propose a plausible molecular mechanism for deafness associated with the mutations at the Ca^2+^ binding residues of the tip-link proteins at the inter-domain linkers. The molecular mechanism deciphered here derives two ways to restore the WT phenomena in mutant tip-links, either by hiking the Ca^2+^ level in endolymph or inducing rigidity to the mutant-linkers with small molecules or osmolytes that can specifically bind to mutant linkers and restore the WT conformation from a bent. The regulator, plasma membrane Ca^2+^-ATPase (PMCA), which regulates the local concentration of Ca^2+^ ions around the tip-links of hair-cells was identified long ago (31), however, our study proposes the use of this molecule as therapeutic to cure certain types of congenital hearing loss. Overall, with the quantitative inputs on the conformations and force-responsive behavior of a zoomed region of the tip-links, we portrayed here the force-propagation pathway through the force-sensors in the inner ear. A continuation in the same line of study would help us to reach the milestone of mapping the force-propagation path in the entire force-sensor complex at the molecular details. A complete force-propagation path would essentially identify all the domains and residues that actively participate in the force-dissipation or propagation and guide us to decipher their contributions in the overall elasticity and the force-conveying behavior. Mutations on the path may have common mechanism of disrupting hearing, and thus a common therapeutic strategy.

## Materials and Methods

### Protein expression and purification

Mouse Cdh23 EC1-2 (Q24 to D228 in NP_075859.2) and Pcdh15 EC1-2 (Q27 to D259 in NP_001136218.1) were recombinantly modified with 6xHis-tag at N-terminal for affinity-purification, and sort-tag (LPETG) at C-terminus for surface attachment (See Table 1 for PCR primer pairs), cloned into Nde1 and Xho1 sites of PET21a vector (Novagen, Merck) and expressed in lemo21 (DE3) (Stratagene) as soluble proteins with 1mM of L-Rhamnose (MP Biomedicals) in the culture media. All the proteins were obtained in cell lysates and subsequently purified using Ni-NTA (Qiagen) affinity-columns and superdex 200 increase 10/300 GL (GE Lifesciences) size-exclusion columns. The purity of the proteins was monitored from the absorbance-based chromatograms, SDS-PAGE gel electrophoresis (Figure S2). Folding was verified using circular dichroism, and the activity of the proteins was checked from the protein-protein interactions using single-molecule pull-down (Figure S3).

### Labeling of fluorophores on proteins

For attaching Cy3 to the N-terminal of Cdh23, Cdh23(V2C) was reacted with Cy3 maleimide at a protein:dye ratio of 1:8 at room temperature for 8 hours. The unreacted fluorophores were separated using 10 kDa spin columns, and the final labeling ratio of protein:dye was maintained to 1:1. Cy5 was attached to the C-termini of Pcdh15 constructs using sortagging. The proteins were recombinantly modified with sortase A recognition tag (LPETGGSS) at the C-termini. Sortase A enzyme, recognizes this sequence and form a thioester intermediate, which is prone towards nucleophilic attack from a polyglycene. We used GGGGC as the nucleophile, and before its use, we modified the polypeptide with Cy5 maleimide (33).

### Single-molecule Fluorescence Resonance Energy Transfer (smFRET) Experiments

The glass-coverslips were cleaned with piranha solution (H_2_SO_4_ : H_2_O_2_ in a ratio of 3:1) for 2 hours followed by plasma treatment for 2 minutes, and finally etched in 1 M of potassium hydroxide. Subsequently, the surfaces were silanized using 2% APTES (Sigma-Aldrich) in Acetone and cured at 110 °C for 1 h. The amine exposed surfaces were reacted with a mixture of 10% NHS-PEG-maleimide (MW: 5kD) in NHS-PEGm (MW:5 kDa) (LaysanBio) in a basic buffer (200mM NaHCO_3_, 600mM K_2_SO_4_, pH 8.6) for 4 hours. The PEG-modified surfaces were then incubated in 100 μM polyglycine (GGGC) for 7 h at 25 °C to anchor the peptide to surfaces via cysteine-maleimide reaction. This polyglycine serves as a nucleophile for sortagging. Next, we used the sortagging protocol as discussed previously to anchor the C-terminal of Cy3 labeled Cdh23 (WT) onto the polyglycene coated surfaces where the N-terminal of the polyglycene serves as a nucleophile (33). To achieve distributed fluorescence spots on the surface, we mixed the labeled and unlabeled protein in 1:10. Next, we monitored the extent of surface functionalization using a total internal reflection fluorescence microscope (IX83 P2ZF inverted microscope equipped with IX83 MITICO TIRF illuminator). We excited the surface at 532 nm laser at an exposure time of 300 ms. Fluorescence was collected using 60X oil immersion objective and split into four channels according to wavelength using quad-band filter recorded using EMCCD coupled camera. For a good and acceptable surface, we observed discrete single molecule signals only at the cy3 channel.

Next, we incubated the surface with 50 nM Cy5-labeled Pcdh15 for an hour, followed by an extensive wash with pH 7 buffer, and monitored FRET with 532 nm laser at an exposure time of 300 ms. smFRET was performed at room temperature in 25 mM HEPES buffer containing 50 mM KCl, 50 mM NaCl, 2 mM Trolox. FRET efficiency was calculated using the equation,

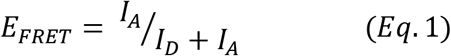

Here, *I_A_* and *I_D_* are background corrected, and leakage corrected fluorescence intensity of donor and acceptor respectively.

For ensemble FRET at varying calcium concentrations, we first chelated out calcium from proteins using chelex resin. For experiments, an equimolar amount of Cdh23(WT) and Pcdh15(WT) were mixed first and then titrated with calcium stepwise while recording fluorescence using steady-state fluorescence spectrophotometer (Horiba Jobin Yvon). FRET was monitored by exciting the solution at 545 nm wavelength and monitoring the emission from 555-720 nm wavelength maintaining excitation and emission slits of 3 and 5 nm respectively.

### Single-Molecule Force spectroscopy using AFM

For single molecule force-spectroscopy using AFM (Nano wizard 3, JPK Instruments, Germany), the C-termini of the protein molecules were immobilized on the freshly cleaned glass-coverslips and Si_3_N_4_ cantilevers (Olympus, OMCL-TR400PSA-1), using sortagging, as described previously in the context of smFRET (31). To achieve a higher density of molecules on the surface, we used 10% bifunctional PEG (NHS-PEG-maleimide, MW 5000Da) in mono-functional PEG (NHS-PEG, MW 5000Da). With this mixture, we achieved a separation between two neighboring molecules in ∼150 nm (33).

For dynamic force-ramp measurements, we brought the cantilever down in contact at 2000 nm.s^−1^, waited for 0.5 s for proteins to interact, and finally retracted at velocities varying from 200, 500, 1000, 2000, 3000 and 5000 nm.s^−1^. At each pulling velocity, we performed 5625 force-curves at five different positions. We repeated the experiments three times with different batches of proteins and merged all data. We always measured the spring constant of the cantilevers before and after each experiment from the power spectra of the thermal oscillations. The spring constant of the modified cantilevers falls in the range of 20-30 pN.nm^−1^.

To distinguish the specific PEG-stretching from non-specific events, we fitted the force-curves to the following freely-jointed chain model (FJC) (55)

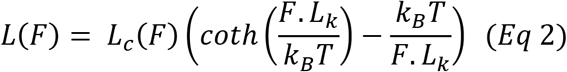

Where *L_k_* is the Kuhn length, and *L_c_* is the contour length of PEG, *k_B_* is Boltzman constantand T is the temperature. The unbinding force was measured from peak maxima of each force curve and plotted as Force histograms for different pulling velocities. The loading rate was measured using two widely used methods. In one case we measured directly from spring constant and velocity which defines one loading rate/velocity. Secondly, we followed the model given by Ray et al. for estimathe tion of loading rate from each force-extension profile of each pulling velocities (37). We used the following equation

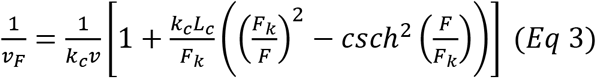

where *L_c_* is the contour length, *k_c_* is the spring constant of the cantilever, *v* is the velocity of the cantilever, *a* is the kuhn length, and 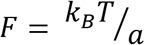.

Along with, we also monitored the distribution of contour lengths (L_c_).

The most probable force (*F*) with loading-rate (*v_F_*) was fitted to Bell-Evans model and estimated the kinetic parameters like off-rate (*k*_0*ff*_), transition distance (*x_β_*) using the following equation:

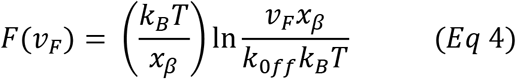

### Molecular Dynamics Simulations

All the molecular dynamics (MD) simulations were set up using the QwikMD (45)plugin of VMD (46) and run using NAMD (47) version 2.12. Available crystal structures were used as start-structures for simulations. For PCDH15-D157G, where structural data was not available, we made the mutation using QwikMD from the WT-structure. The rest of the setup and simulation protocol was identical for all the complexes. After the addition of hydrogen atoms, the structures were aligned with the longest axis along the Z-axis and solvated with TIP3P water to form a box with a distance of 15 Å between the edge of the box and the protein in all the axis. The Na^+^ and Cl^−^ ions corresponding to a concentration of 150 mM were then placed randomly in the water box by replacing the water molecules. The system was then minimized for 5000 steps with the position of the protein atoms fixed followed by 5000 steps without any restraints. The temperature of the system was then gradually raised to 300K at a rate of 1 K every 600 steps with the backbone atoms restrained. This was followed by equilibration for 5 ns with the backbone atoms restrained followed by 10 ns of the production run with no restraints. For all the steps, the pressure was maintained at 1 atm using Nose-Hoover method (56, 57), thelong-range interactions were treated using the particle-mesh Ewald (PME) (57) method and the equations of motion were integrated using the r-RESPA scheme to update short-range van der Waals interactions every step and long-range electrostatic interactions every 2 steps. The MD refined structures were used for Steered Molecular Dynamics (SMD) simulations after addition of extra water towards the pulled end and minimization, heating and equilibration of the added water with constraints on the protein backbone. The C terminus of Pcdh15 was fixed,and C-terminus of Cdh23 was pulled with constant velocity ranging from 1 Å/ns, 2.5 Å/ns, 5 Å/ns and 7.5 Å/ns.

### Dynamic Network Analysis

The Dynamic Network Analysis was performed using the VMD’s Network View Plugin (49) and the associated scripts. The network was created by considering all the α-carbons as nodes and creating a connecting edge between them if the two residues had a heavy (non-hydrogen) atom within a distance of 4.5 Å from each other for 75% of the trajectory. The edges were then weighted using the correlation matrix calculated using the program carma (54). The strength of the correlation between the two nodes determines the probability of information transfer, in this case, force propagation, between them by the relation w_ij_ = −log (|c_ij_|). The neighboring α-carbons in the sequence were ignored as they lead to trivial paths (50). The network was then partitioned into sub-networks or communities using Girvan-Newman algorithm (58). The suboptimal paths up to 20 edges longer than the shortest path were calculated between residue 208 of CDH23 and residue 230 of PCDH15 using Floyd-Warshall algorithm (59). The flexible residues at the ends were not considered in the Network Analysis procedure.

## ASSOCIATED CONTENT

### Supporting Information

Supplementary figures and tables are attached separately.

## Acknowledgments

This work was supported by the Wellcome Trust/DBT India Alliance Fellowship [grant number: IA/I/15/1/501817] awarded to SR.

S.R. acknowledges the financial support by the Wellcome Trust/DBT Intermediate fellowship by India Alliance and Indian Institute of Science Education and Research Mohali (IISERM). JPH, AS, DD sincerely thank IISERM for financial support. NA thanks CSIR-India for fellowships. AS is thankful to the Centre of Excellence (COE) in Frontier Areas of Science and Technology (FAST) program of the Ministry of Human Resource Development, Government of India for financial support.

