## Supplementary for "Broken force dispersal network in tip-links by the mutations induces hearing-loss"

| | | |
|---|---|---|
| Figure S1: | Determination of calcium binding constant using competitive Chelator method | 3 |
| Figure S2: | SDS-PAGE gel picture, Circular Dichroism spectra and size exclusion Chromatography result of the proteins | 4 |
| Figure S3: | Single-molecule pull down of proteins using TIRF | 5 |
| Figure S4: | TIRF images of Single-molecule FRET | 6 |
| Figure S5: | Characteristic single molecule PEG stretching curve | 7 |
| Figure S6: | PEG extension profile for various protein pairs | 8 |
| Figure S7: | Percentage of single molecule events obtained and contour length distribution | 9 |
| Figure S8: | Force histogram of mutant complex at 2mM $Ca^{2+}$ buffer | 10 |
| Figure S9: | Bio-layer interferometry result for the interaction | 11 |
| Figure S10: | Cartesian principal compoenntanalsyis result of Molecular dynamics simulation of heteromeric wild-type and mutant complexes | 12 |
| Figure S11: | H-bond analysis of MD simualtionperfomed for various complexes | 13 |
| Figure S12: | Averaged strcture of the most populated mutant conformation during the Simulaiton | 14 |
| Figure S13: | Angle of force propagation vectors with pulling vector | 15 |
| Figure S14: | Cirtical nodes and edges of force propagation | 16 |
| Text S1. | Competitive chelator method for calcium binding constant determination | 17 |
| Table S1: | Cdh23 (WT) and Cdh23 (D101G) $Ca^{2+}$ dissociation constants | 18 |
| Table S2: | Pcdh15 (WT) and Pcdh15 (D157G) $Ca^{2+}$ dissociation constants | 18 |
| Table S3: | FRET efficiency and FRET distances for various complexes at varying $Ca^{2+}$ concentrations | 19 |
| Table S4: | Single-molecule event rate obtained at different pulling velocities | 20 |
| Table S5: | Off-rate and on-rate obtained from BLI measurements | 20 |
| Table S6: | Experimentally calculated loading rates for each pulling speed | 21 |

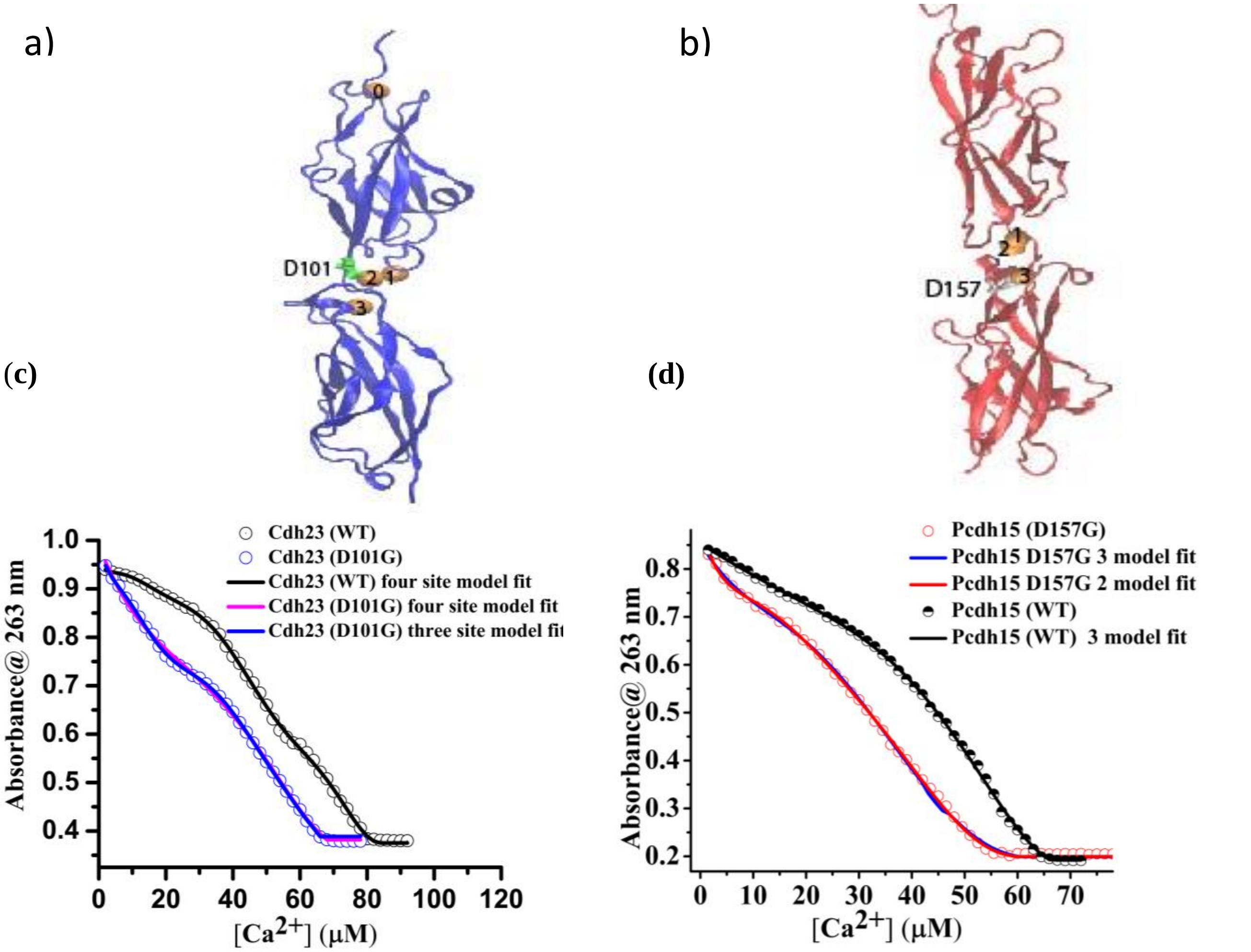


**Figure S1: Determination of calcium binding constant using competitive chelator method (In support of Figure 1)** (a) Structure of Cdh23(WT) marking the calcium ions. Aspartate at 101th position has been shown as green stick connected to 2nd calcium ion (b) Structure of Pcdh15(WT) with calcium ions (marked with number). Aspartate at 157th position has been showed in stick and connected to 3rd calcium ion (c) Gradual decrease in the absorbance of BAPTA at 263 nm with increasing $Ca^{2+}$ concentration was monitored in presence of 25 μM of Cdh23 EC1-2(WT) (Black Circle), and in 25 μM of Cdh23 EC1-2(D101G) (Blue circles) independently (for further experimental procedures see text S1). Black line represents the four sites binding model fit for the WT. Blue solid line represents the fitting using three sites binding model and magenta line represents fitting using four sites binding model for the data correspond to Cdh23 EC1-2(D101G). The four binding site model for WT and three model fit for mutant was accepted from the F-test. (c) Gradual decrease in the absorbance of BAPTA at 263 nm with increasing $Ca^{2+}$ concentration was monitored in presence of 25 μM of Pcdh15 EC1-2(WT) (Black Circle), and in 25 μM of Pcdh15 EC1-2(D157G) (red circles) independently. Black line represents the fitted curve obtained using three sites binding model for WT. Red and blue line represent the fitted curves obtained from three and two binding sites models respectively for Pcdh15 EC1-2(D157G). Three binding model was accepted for both WT and mutant from the F-test.

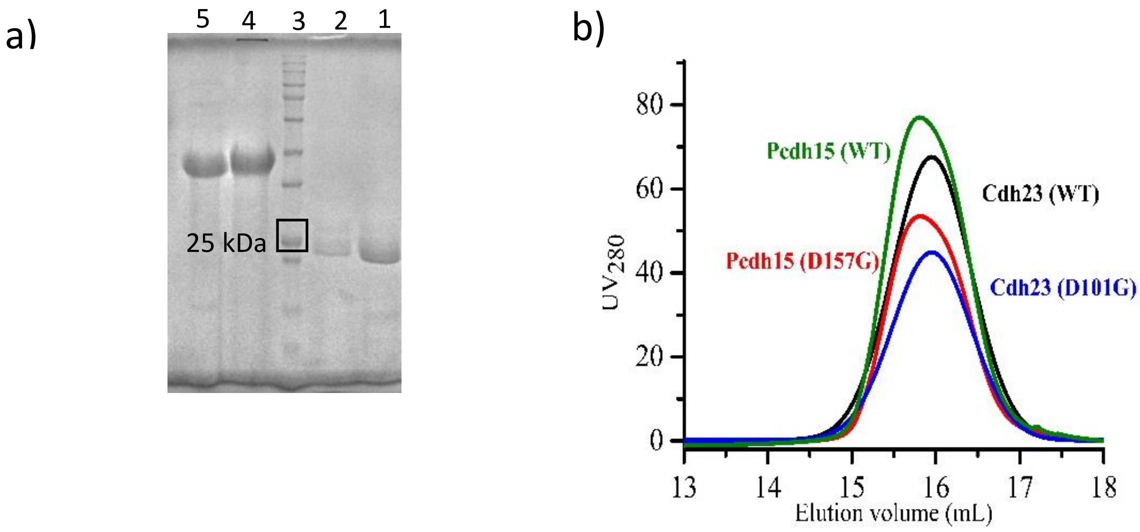


1- Cdh23 (WT)
2- Cdh23 (D101G)
3- Protein molecular weight marker
4- Pcdh15 (WT)
5- Pcdh15 (D157G)

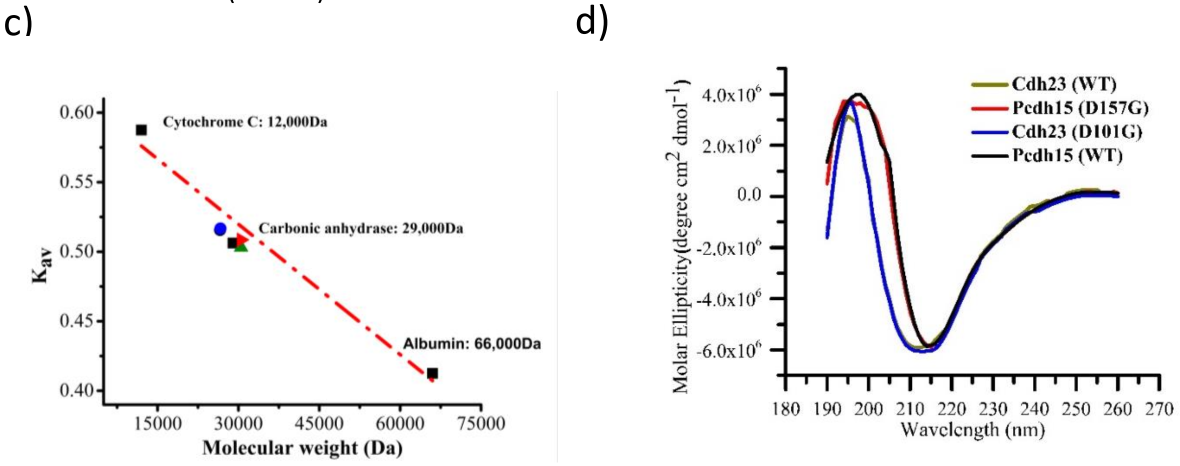


**Figure S2: Purity and folding of the proteins** (a) SDS-PAGE picture of all protein variants after size exclusion chromatography (b) Size exclusion chromatography (SEC) profile of the proteins performed at 50 µM $Ca^{2+}$ using S200 GL column. (c) Putting the elution volumes in the calibration curve suggests the monomeric elution for all proteins in the monomeric forms. (d) Circular dichroism spectra of all the proteins obtained at 50 µM $Ca^{2+}$ buffer showing the peak at 216 nm denoting the β -sheet rich structure of all the proteins.

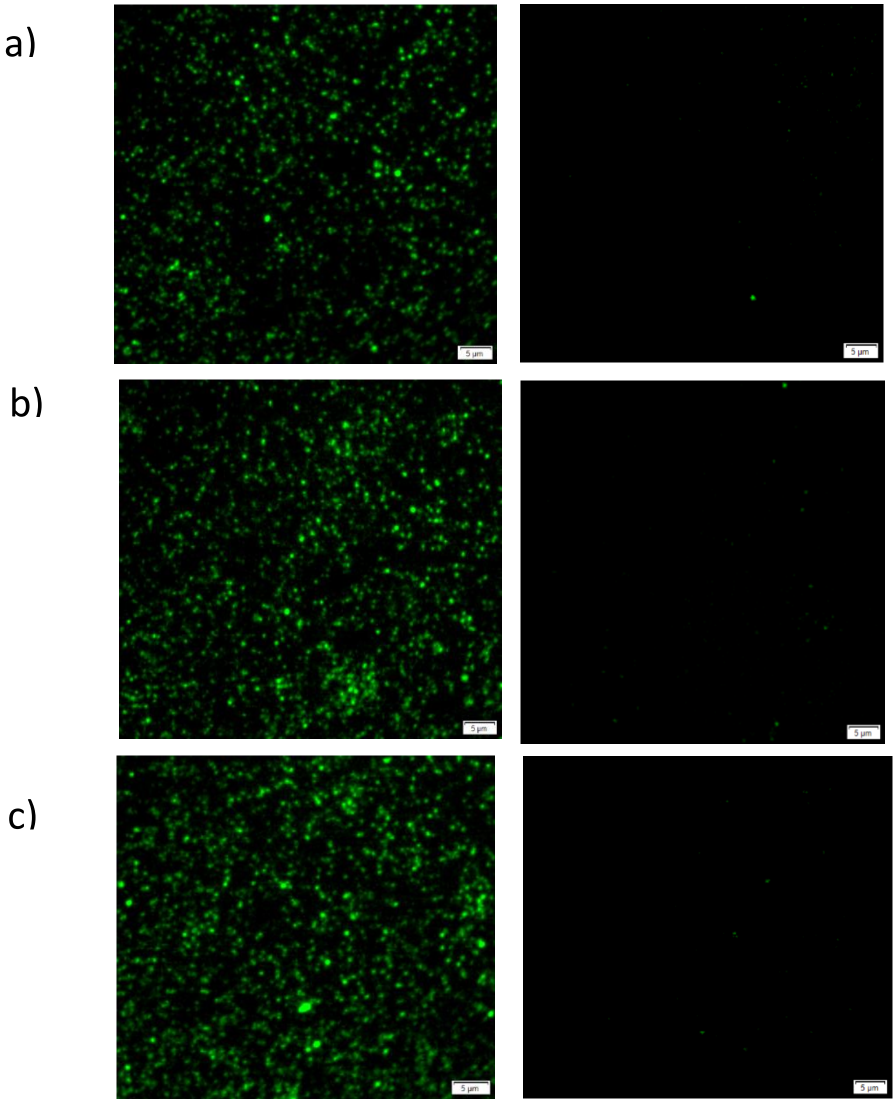


**Figure S3: Single-molecule pull-down images depicting the functionality of the protein(In support of Figure 1) (**a) We covalently immobilized the C-terminal of unlabelled Cdh23(WT) on a glass surface. Next, we incubated the surface with Cy3 labeled Pcdh15(WT) for 15 mins followed by thorough washing. We then visualized the surface under TIRF microscope by exciting with 532 nm laser at an exposure time of 300 ms and observed fluorescence signals (green spots) from the Cy3-Pcdh15(WT) pulled down by Cdh23(WT). To monitor the specificity, we incubated the surface with 5 mM EGTA buffer for half an hour under dark where EGTA chelates all $Ca^{2+}$ out and monitored the signal again (right panel in a). We observed 92% drop in fluorescence signal, suggesting the interaction is $Ca^{2+}$ dependent. (b) The pull-down of Cy3- Cdh23(D101G) by surface attached Pcdh15 EC1-2(WT) was performed, and subsequent loss of signal was monitored after EGTA wash (Right panel). (c) The pull-down of Cy3- PCdh15 (D157G) by surface attached Cdh23 EC1-2(WT) was performed, and subsequent loss of signal was monitored after EGTA wash (Right panel).

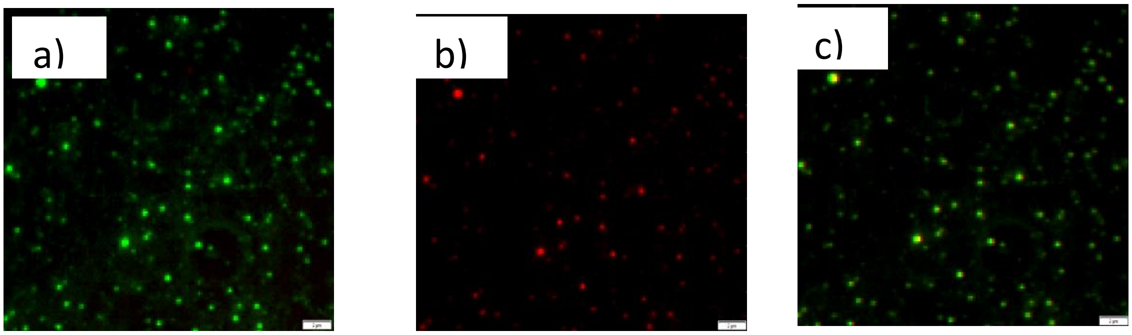


**Figure S4: Representative TIRF images for the smFRET performed with Cy3-Cdh23 EC1-2 on the surface incubated with Cy5-labeled Pcdh15 (In support of Figure 1)** (a) Signal at the donor Cy3 channel, and (b) at the acceptor Cy5 channel. (c) Overlay of the Cy3 and Cy5 channel depicting the co-localization of the spots of both the channels. These images are representative of all three pairs, Cdh23(WT): Pcdh15(WT); Cdh23(WT):Pcdh15(D157G); and Cdh23(D101G):Pcdh15(WT), at varying $Ca^{2+}$.

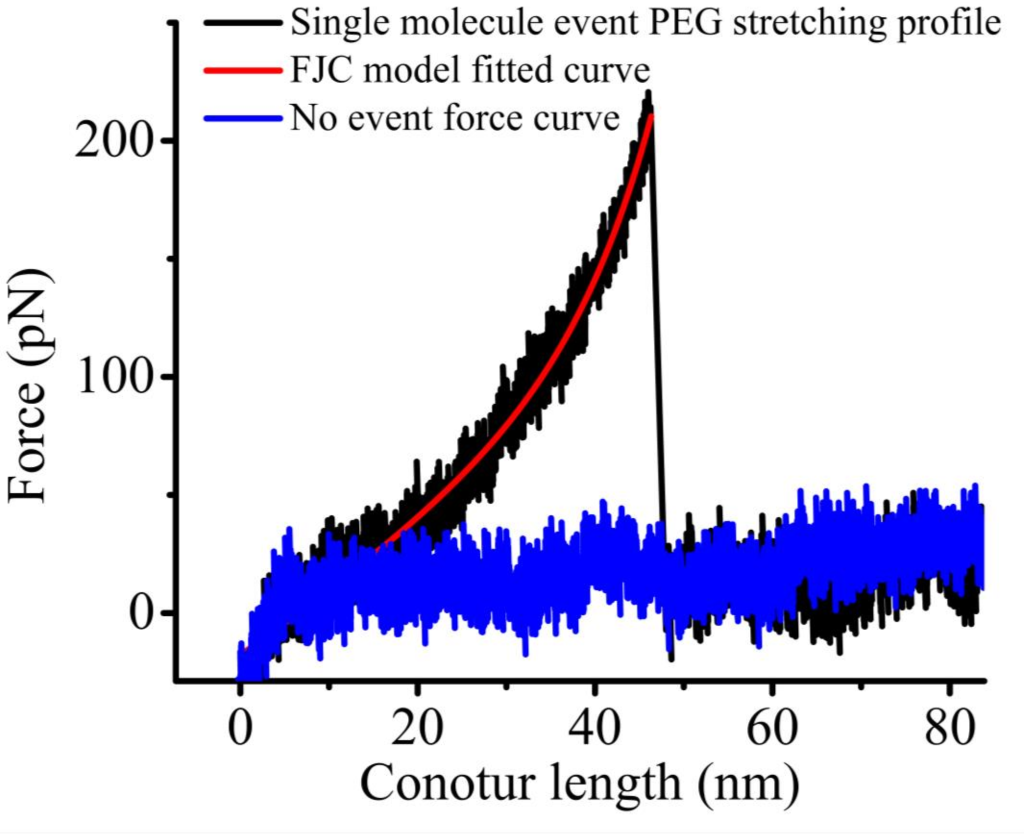


**Figure S5: Characteristic single-molecule PEG stretching event with FJC model fit (In support of Figure 2)** Black curve shows typical PEG stretching obtained during a single molecule force spectroscopy event along with FJC model fitting (red). Blue line depicts typical curve profile when there is no event.

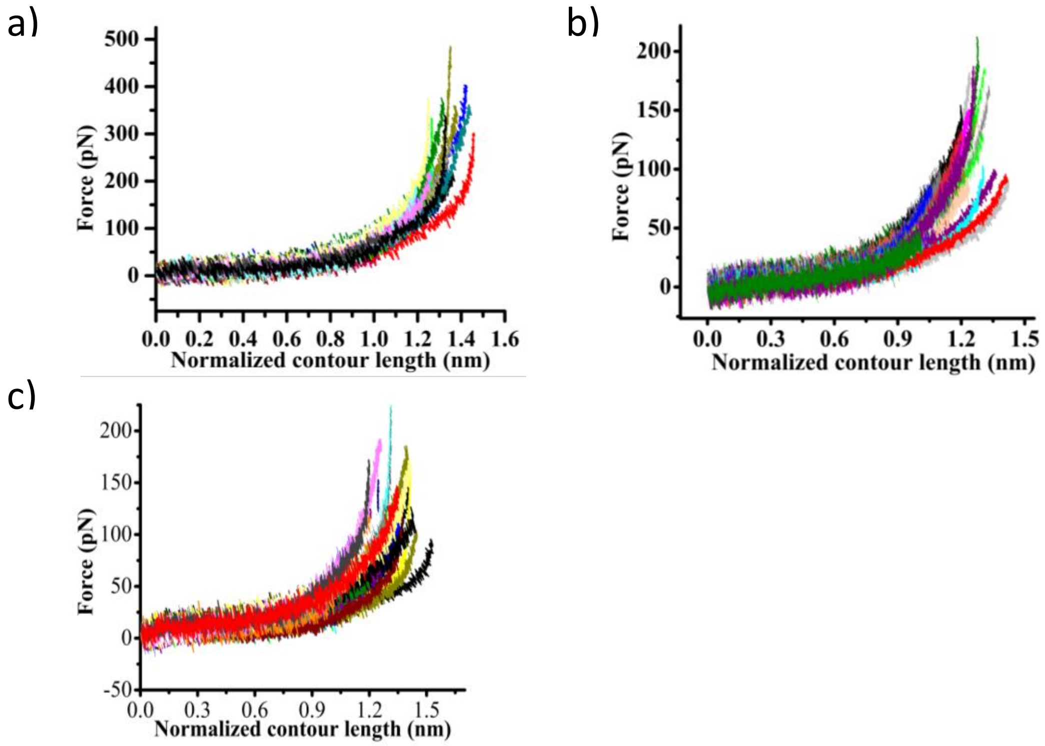


**Figure S6. Selected experimental force-stretching curves for different protein pairs at pulling speed of 2000 $nm.s^{-1}$ (In support of figure 2)** (a) Cdh23 EC1-2 (WT) vs. Pcdh15 EC1-2(WT) interactions; (b) Cdh23 EC1-2 (D101G) vs. Pcdh15 EC1-2(WT) interactions; and (a) Cdh23 EC1-2 (WT) vs. Pcdh15 EC1-2(D157G) interactions. The curves are normalized by the extension (nm) corresponding to a constant force of 100 pN. The normalization protocol is described previously[13,14].

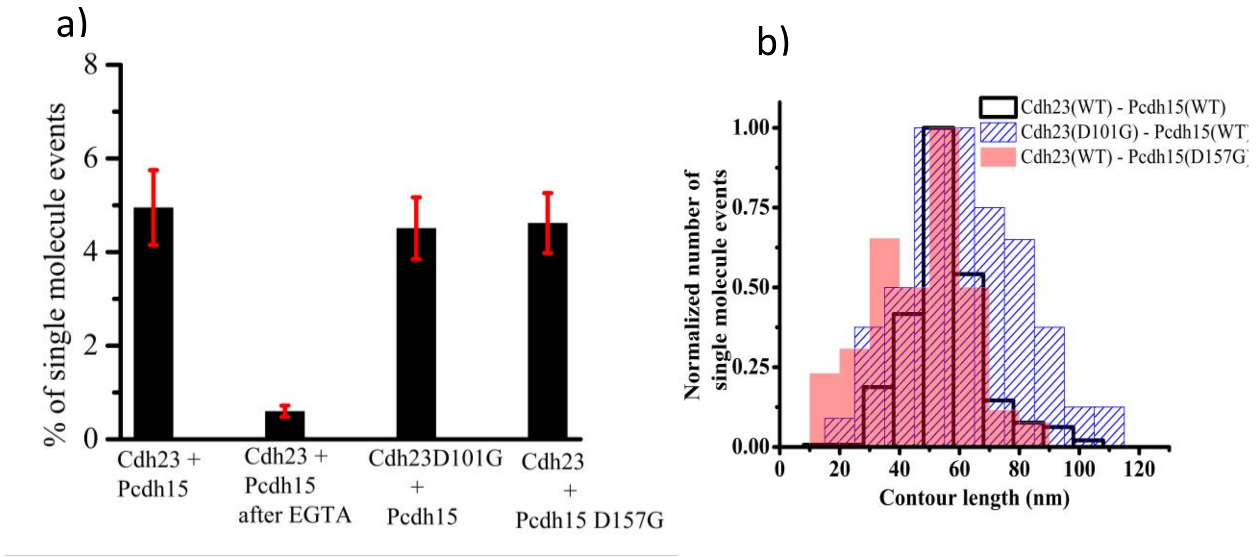


**Figure S7: Percentage of single-molecule force-stretching events and the distribution of contour length of PEG (In support of Figure 2)**(a) Percentage of single-molecule stretching events obtained for different protein pairs in presence and absence of $Ca^{2+}$ are shown. (b) Contour-length of PEG is obtained from the fitting of each PEG stretching to WLC model as shown in S6. The distributions of contour-length for WT Cdh23 – WT Pcdh15 (black), Cdh23 D101G – WT Pcdh15 (blue) and WT Cdh23 – Pcdh15 D157G (red shaded) are similar and peaked at the theoretical estimate of two PEG stretching in series.

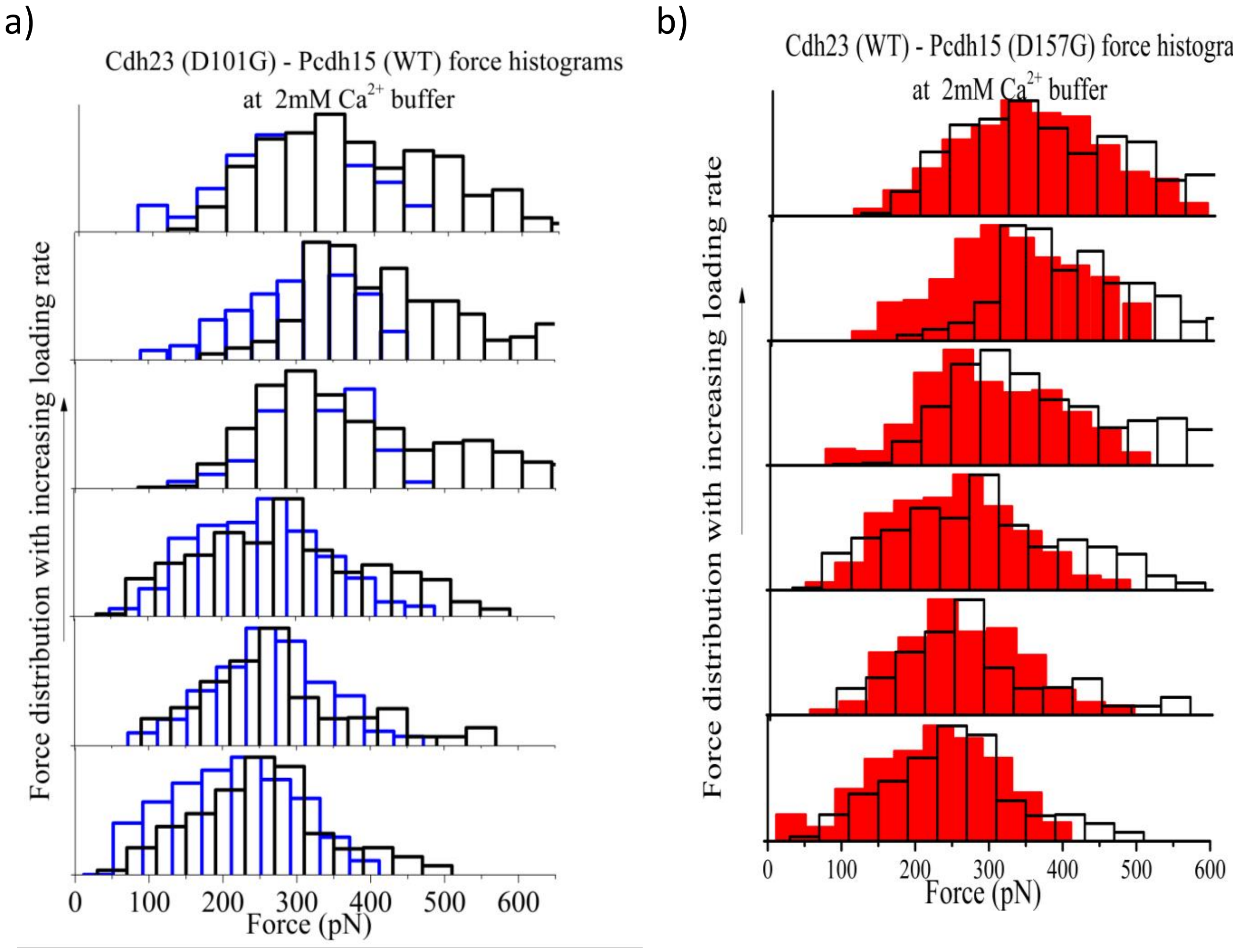


**Figure S8: Force histogram of WT-mutant pairs at 2mM $Ca^{2+}$ buffer. (In support of figure 2)**(a) Force distributions obtained at different pulling velocities (200, 500, 10000, 2000, 30000, 50000 $nms^{-1}$) for Cdh23 (WT) – Pcdh15(WT) complexes (black) overlapped with Cdh23(D101G) – Pcdh15(WT) complex (blue). It shows that the force bearing property of the mutant complex gets restored at elevated calcium level. (b) Force distributions obtained at different pulling velocities (200, 500, 10000, 2000, 30000, 50000 $nms^{-1}$) for Cdh23 (WT) – Pcdh15 (WT) complexes (black) overlapped with Cdh23 (WT) – Pcdh15 (D157G) complex (red shaded). For this mutant also force sensing property becomes comparable to the WT complex at high calcium cocnentration.

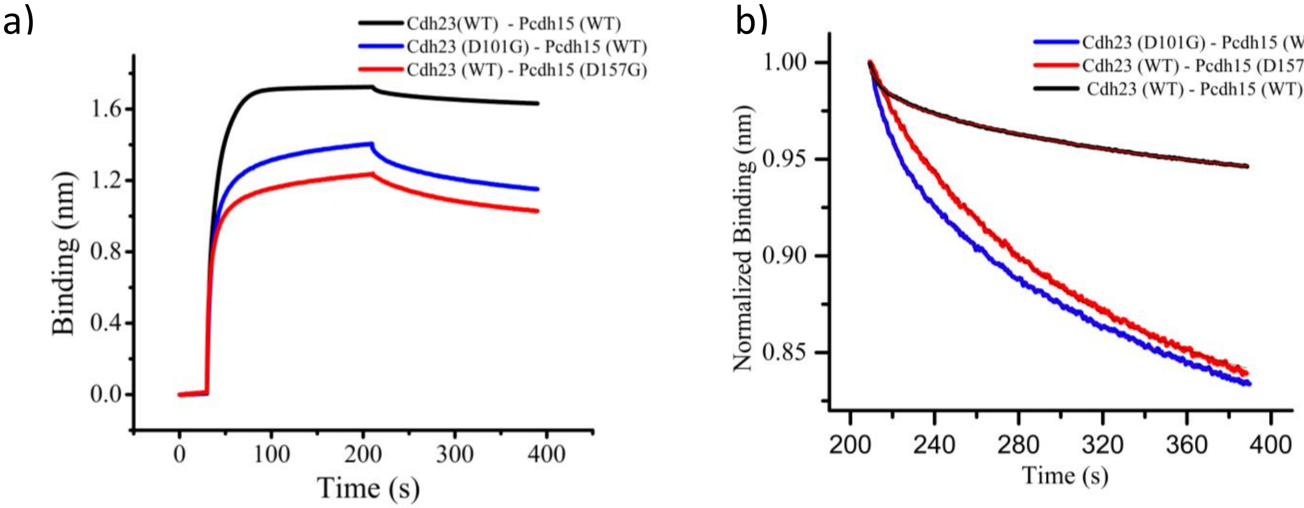


**Figure S9: Bio-layer interferometry (BLI) to monitor the interactions between Cdh23(WT) Pcdh15(WT) and mutnat complexes (In support of Figure 2)** We carried out Bio-layer interferometry (BLI) measurements using Amino-Propyl Silane (APS) sensor tip (Pall life science). We modified the APS tip with bifunctional NHS-PEG-Maleimide followed by polyglycine attachment as described above for smFRET and force spectroscopy. The density of the bifunctional PEG was maintained at 100 %. Next, the C-terminal of one of the protomers (analyte) was attached to the tip using sortagging, and the other protein was used as a ligand in 1 mL tube. Each cycle in the experiment consists of three steps. Step 1 is the baseline-correction where the sensor tip is immersed in the buffer alone for 30 s. Step II is the association step where the tip is dipped into the ligand proteins for 300 s,. The third step is the dissociation step where the tip is again immersed in the buffer for 300 s. Data-fitting to 1:1 reaction model was done using the inbuilt fitting program of the Instrument. (a) The complete binding run for Cdh23(WT) - Pcdh15(WT), Cdh23(D101G) – Pcdh15(WT), and Cdh23(WT)-Pcdh15 (D157G) performed at 50 μM $Ca^{2+}$ buffer are shown here. This depicts that the mutants are having lower on-rate and higher off-rate. (b) A normalized plot of the binding only during dissociation is highlighted here which clearly demostrates the lower off-rates for mutant-complexes. The rates were estimated from the 1:1 reaction modelfit to the data (mentioned in main-text).

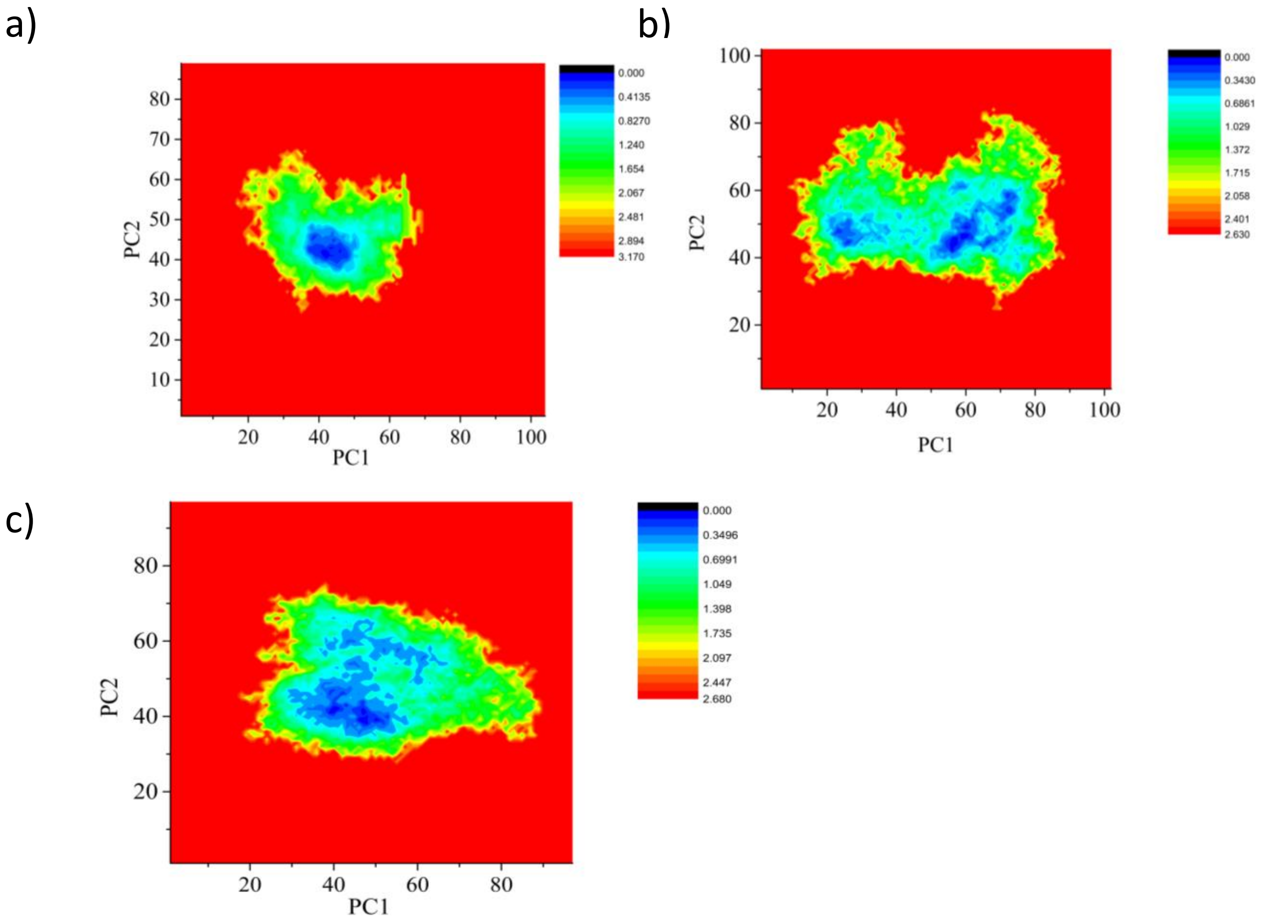


**Figure S10: PCA based Free Energy landscape for the WT and mutant complexes (In support of Figure 3).** The conformations visited during the three trajectories are projected on the Free Energy Landscape using the 1st and 2nd principal components as the reaction coordinates.(a) WT complex (with all $Ca^{2+}$), (b) Cdh23 (WT)-Pcdh15 (D157G) complex (3rd $Ca^{2+}$-removed), and (c) Cdh23 (D101G)-Pcdh15 (WT) mutant complex (2nd $Ca^{2+}$-removed). It is observed that the energy landscape of the WT complex has one well-defined energy basin, which is occupied by most of the conformations. However, the mutants have much higher conformational heterogeneity leading to broader occupancy of the energy landscape and the conformations are concentrated in two major energy basins with multiple small clusters inside them. This indicates that WT complex adopts a relatively rigid conformation but the complexes of the mutants are more flexible and exist is atleast two major conformations.

a) b) c)

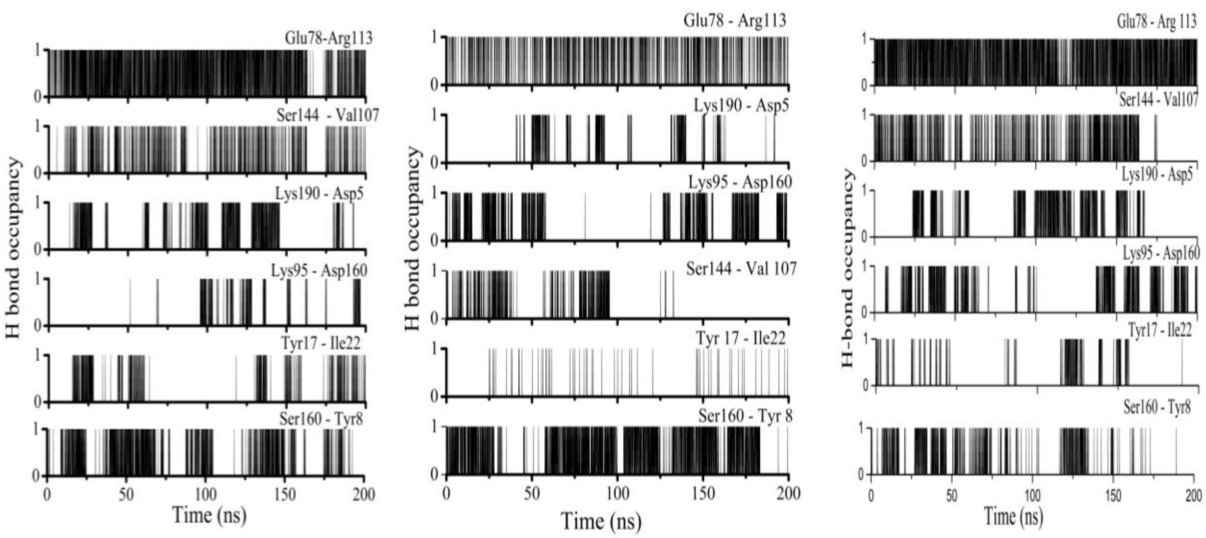


**Figure S11: H-bond analysis of MD simulation (In support of figure 3)** (a) Occupancy of various interprotein (between Cdh23 and Pcdh15) H-bond pairs during gaussian accelerated MD simulation of Cdh23 (WT) – Pcdh15 (WT) complex for 200 ns. (b) Occupancy of various interprotein H-bond pairs during gaussian accelerated MD simulation of Cdh23 (WT) – Pcdh15 (D157G) complex with one calcium ion removed for 200 ns. (c) Interprotein H-bond plots for Cdh23 (D101G) – Pcdh15 (WT)Complex simulation with one calcium removed performed for 200 ns.

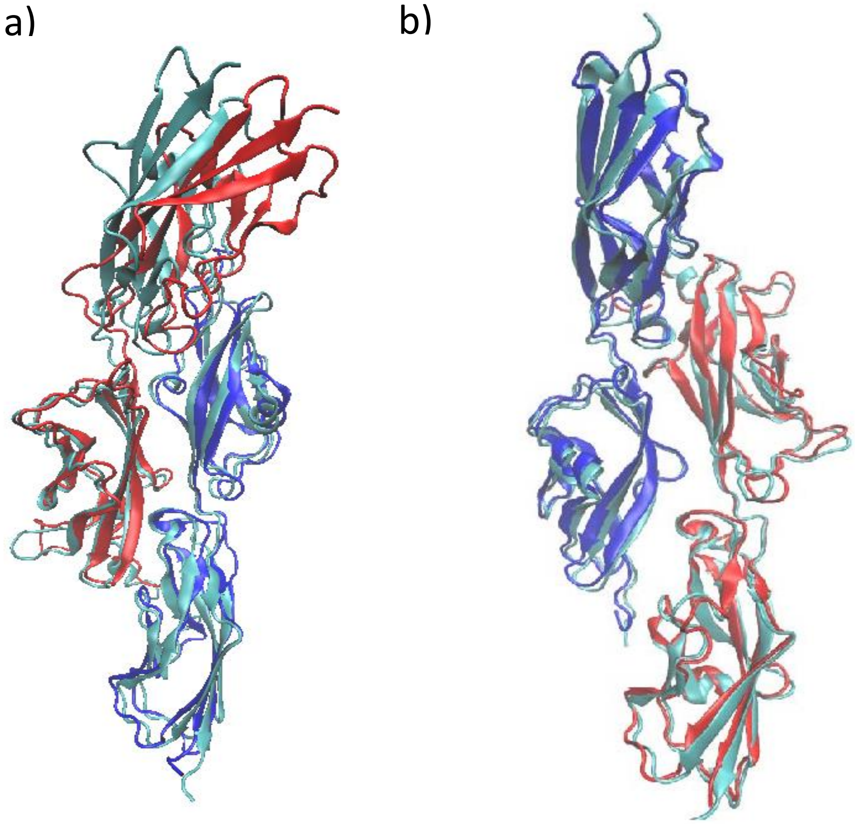


**Figure S12: The most probable conformation of mutant complexes are overlapped with WT complex as obtained from cPCA (In support of figure 3)**The most probable conformations were derived by averaging all the conformations obtained from the most populated cluster in the cPCA. (a) Overlap of WT with mutant-1 (Cdh23 (WT) – Pcdh15 (D157G) (red and blue, red denoting Pcdh15 and blue denoting Cdh23) and (b). Overlap of WT with mutant-2 (Cdh23 (D101G) – Pcdh15 (WT) (red and blue, red denoting Pcdh15 and blue denoting Cdh23). WT complex is denoted as cyan in both cases.

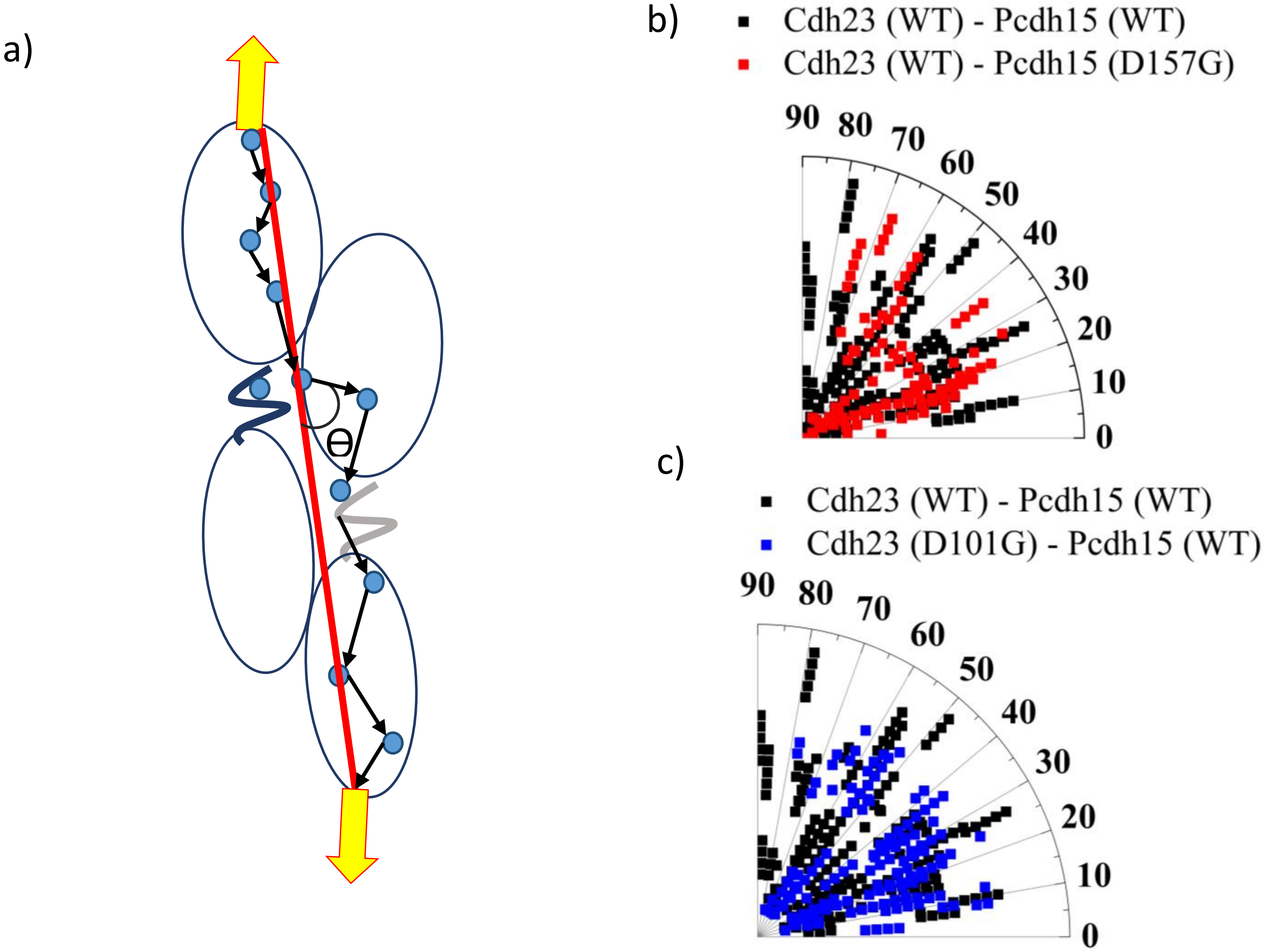


**Figure S13: Distributions in the angle of force propagation vectors with the direction of pulling**. **(In support of figure 3)** (a) Schematic description of the direction of pulling in SMD along with the angles measured between the edges connecting nodes (vectors, black arrows) and the pulling axis (red line) (Methods) for each suboptimal path. The black arrows describe the force propagation from one node to the next one. Nodes are represented in blue circles. (b) Comparison of the distributions of the angle of force propagation vectors for WT (black) and mutant-1 (red), and (c) for WT (black) and mutant-2 complex (blue). The vectors make smaller angles for both the mutant complexes than WT, indicating a smaller contribution in the perpendicular component of force-dispersal than the WT complex.

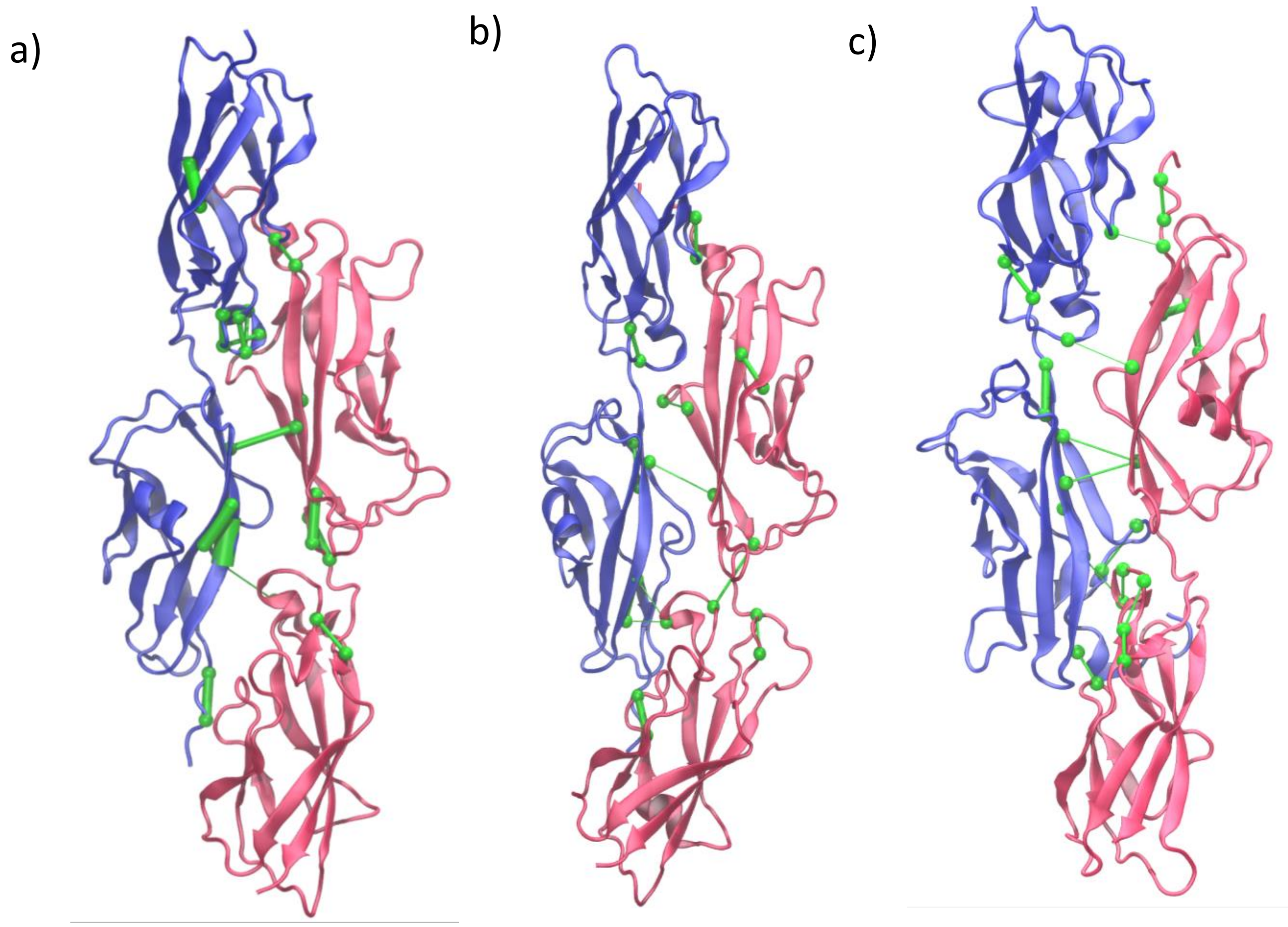


**Figure S14: Depiction of critical nodes and edges along the force propagation pathway for WT and mutant complexes. (In support of figure 3)** (a) Weighted critical nodes and edges for Cdh23 (WT) – Pcdh15 (WT) complex. (b) Cdh23 (WT) – Pcdh15 (D157G) complex. (c) Cdh23 (D101G) – Pcdh15 (WT) complex. Critical nodes were defined on the bases of their 'betweeness'. The correlation in the motion of the nodes was used to weight the edges and is depicted as the width of the edges. The WT complex is connected with more nodes and stronger (thicker) edges compared to D157G and D101G complexes indicating that the force-dispersal in WT complex is more than the mutants, thus making the WT more resilient to force and less vulnerable to force induced dissociation.

**Text S1. Competitive chelator method for calcium binding constant determination**

For the determination of calcium dissociation constant using competitive chelator method we used both quin 2 and BAPTA as calcium indicator. First, a stock 1 mM solution of the indicator was prepared in calcium free buffer. Protein solution was incubated with chelex resin for 2 hours to remove the calcium form the protein. Chelex was removed from the protein solution by centrifuging the solution at 2000 RPM for 10 minutes. Then equimolar amount (25 µM) of protein and indicator were mixed which will be used as titrant. Then calcium was added into the solution in a stepwise manner increasing the concentration at each step 3 µM and at each step absorption spectra was recorded from 200-500 nm wavelength using UV-Visible absorption spectrophotometer (Cary win UV-Visible spectrophotometer). From the spectra absorption change at 262 nm from each step was plotted with calcium concentration after each step. The curve was fitted with the equations 13 and 14 mentioned in reference no. 25 in various models i.e three binding sites, two binding sites, four binding sites model. Fitting was performed using MATLAB program written in-house. Earlier Sotomayor et, al determined the calcium binding affinity of Cdh23 (WT) using fluorescence based competitive chelator method and our results corroborated with their findings.

**Table S1: Estimated $Ca^{2+}$ dissociation constants of Cdh23 EC1-2 (WT) and Cdh23 EC1-2 (D101G) measured using competitive chelator method**

| Cdh23 (WT) (Four-sites model) | Cdh23 (D101G) | |
|---|---|---|
| | Four sites model | three sites model |
| 2.8 ± 0.7<br>8.7 ± 3.2<br>34.2 ± 7.1<br>63.8 ± 11.4 | 3.6± 0.9<br>13.5 ± 3.7<br>99.2 ± 16.5<br>146.7 ± 23.8 | 5.7± 2.6<br>46.2± 9.1<br>98.2± 17.6 |

**Table S2: Estimated $Ca^{2+}$ dissociation constants of Pcdh15 EC1-2 (WT) and Pcdh15 EC1-2 (D157G) measured using competitive chelator method**

| PCdh15 (WT) (Three-sites model) | Pcdh15 (D157G) | |
|---|---|---|
| | Three sites model | Two sites model |
| 34.6 ± 8.8<br>11.2 ± 6.5<br>42.5 ± 9.4 | 44.8± 7.4<br>16.3 ± 4.1<br>152.3 ± 11.9 | 46.5± 8.5<br>20.7± 4.7 |

**Table S3: FRET efficiency and FRET distances for various complexes at different $Ca^{2+}$concentrations (from smFRET experiments)**

| Protein pair | $E_{FRET}$ | Distance (nm) |
|---|---|---|
| Cdh23 (WT) – Pcdh15 (WT) in 50 µM $Ca^{2+}$ buffer | 0.67±0.04 | 4.62±0.02 |
| Cdh23 (WT) – Pcdh15 (WT) in 2 mM $Ca^{2+}$ buffer | 0.68±0.05 | 4.59±0.02 |
| Cdh23 (D101G) – Pcdh15 (WT) in 50 µM $Ca^{2+}$ buffer | 0.40±0.03 | 5.56±0.08 |
| Cdh23 (D101G) – Pcdh15 (WT) in 2 mM $Ca^{2+}$ buffer | 0.66±0.03 | 4.65±0.02 |
| Cdh23 (WT) – Pcdh15 (D157G) in 50 µM $Ca^{2+}$ buffer | 0.48±0.02 | 5.26±0.06 |
| Cdh23 (WT) – Pcdh15 (D157G) in 2 mM $Ca^{2+}$ buffer | 0.67±0.04 | 4.58±0.02 |

**Table S4: Single molecule force-extension events obtained at different pulling velocities.**

| Cantilever Pulling velocity (nm.$s^{-1}$) | Number of events for Cdh23 (WT) – Pcdh15 (WT) out of 5625 measurements (Event-rate in %) | Number of events for Cdh23 (D101G) – Pcdh15 (WT) out of 5625 measurements (Event-rate in %) | Number of events for Cdh23 (WT) – Pcdh15 (D157G) out of 5625 measurements (Event-rate in %) |
|---|---|---|---|
| 200 | 244 (4.3) | 252(4.4) | 277(4.9) |
| 500 | 268 (4.7) | 311(5.5) | 246(4.3) |
| 1000 | 232(4.1) | 216(3.8) | 293(5.2) |
| 2000 | 327(5.8) | 284(5.0) | 213(3.7) |
| 3000 | 287(5.1) | 209(3.7) | 302(5.3) |
| 5000 | 323(5.7) | 268(4.7) | 241(4.2) |

**Table S5: Off-rate and on-rate for protein complexes obtained from BLI measurements**

| Protein pair | Off-rate ($s^{-1}$) | On rate ($M^{-1}s^{-1}$) |
|---|---|---|
| Cdh23 (WT) – Pcdh15 (WT) | 6.6E-04 ± 2.8E-05 | 5.1E+04 ± 2.4E+02 |
| Cdh23 (D101G) – Pcdh15 (WT) | 5.8E-03 ± 7.4E-04 | 9.9E+03 ± 5.9E+01 |
| Cdh23 (WT) – Pcdh15 (D157G) | 3.2E-03 ± 6.6E-04 | 6.5E+03 ± 2.3E+01 |

**Table S6: Loading rates estimeated from each force-extension curves using model given by Ray et al[2] for all pairs of interaction**

| | Cdh23 (WT) – Pcdh15 (WT) | | Cdh23 (D101G) – Pcdh15 (WT) | | Cdh23 (WT) – Pcdh15 (D157G) | |
|---|---|---|---|---|---|---|
| Pulling velocity | Loading rate ($pN.s^{-1}$) | Standard error of mean | Loading rate ($pN.s^{-1}$) | Standard error of mean | Loading rate ($pN.s^{-1}$) | Standard error of mean |
| 200 | 6632 | 56 | 5126 | 32 | 4165 | 40 |
| 500 | 16427 | 420 | 10478 | 52 | 11872 | 360 |
| 1000 | 38244 | 459 | 31024 | 120 | 36318 | 286 |
| 2000 | 58703 | 622 | 48211 | 223 | 47022 | 381 |
| 3000 | 92298 | 754 | 88204 | 475 | 81355 | 719 |
| 5000 | 147225 | 1419 | 112540 | 742 | 102114 | 833 |
